# Overturning circulation structures the microbial functional landscape of the South Pacific

**DOI:** 10.1101/2025.01.12.631543

**Authors:** Bethany C. Kolody, Rohan Sachdeva, Hong Zheng, Zoltán Füssy, Eunice Tsang, Rolf E. Sonnerup, Sarah G. Purkey, Eric E. Allen, Jillian F. Banfield, Andrew E. Allen

## Abstract

Global overturning circulation partitions the deep ocean into regions with unique physicochemical characteristics, but the extent to which these water masses represent distinct ecosystems remains unknown. Here, we integrate extensive genomic information with hydrography and water mass age to delineate microbial taxonomic and functional boundaries across the South Pacific. Prokaryotic richness steeply increases with depth in the surface ocean, forming a “phylocline”, below which richness is consistently high, dipping slightly in highly aged water. Reconstructed genomes self-organize into six spatially-distinct taxonomic cohorts and ten functionally-distinct biomes that are primarily structured by wind-driven circulation at the surface and density-driven circulation at depth. Overall, water physicochemistry, modulated at depth by water age, drives microbial diversity and functional potential in the pelagic ocean.

## Main Text

Microbial plankton are fundamental to the biogeochemical cycling of nutrients (1) and organic carbon in the ocean (2). Extensive sampling efforts have expanded our knowledge of the diversity and metabolic potential of pelagic microbial communities, beginning with the Sargasso Sea metagenome (2003)(*3*) and becoming increasingly sophisticated and depth-integrated with the Sorcerer II’s Global Ocean Sampling expedition (*4*–*6*) (2003-2007), *Tara* Oceans (*7, 8*) (2009-2013) and the 2010 Malaspina Expedition (*9*–*11*). More recently, molecular sampling has begun to be incorporated into existing efforts to characterize oceanic geochemistry, such as bioGEOTRACES (*12, 13*) and bio-GO-SHIP (*14*). Despite the sizable volume of molecular data collected, we still lack a fundamental understanding of whether physical forcing and overturning circulation structure oceanic biomes across latitude, depth, temperature, and nutrient regimes.

Beneath the surface mixed layer, the ocean is stratified into water masses of different densities (*15*). Deep ocean water circulates around the planet on the timescale of hundreds of years (*16*), driven by differences in temperature and salinity. This Global Overturning Circulation (GOC) carries microbial plankton through a range of temperature and pressure extremes that would be fatal to most metazoan life (*17*). Water mass properties like temperature, salinity, nutrients, and oxygen concentration are known to structure microbial communities in the surface ocean (*18*– *21*). However, the extent to which water masses and water aging structure populations of microbial plankton, which often turnover on the order of days (*22, 23*) and have varying resilience to temperature (*24*) and pressure (*25*) extremes, is largely unknown. The microbial functional response to GOC has implications for global carbon cycling at all stages, including carbon export, remineralization, and deep ocean storage as recalcitrant dissolved organic matter (*26, 27*).

Here, we report on the species-level biogeography and metabolic potential of pelagic microbes in the South Pacific by analyzing molecular samples collected every half degree of latitude across full ocean depth as part of the Global Ocean Ship-Based Hydrographic Investigations Program (GO-SHIP) (*28*) P18 line (Fig. 1A). We establish cohorts of co-occurring microbes and correlate the resulting spatial distributions with fine-scale physicochemical metadata to build a bottom-up understanding of the factors driving microbial biogeography. We leverage high-resolution temperature, salinity, and nutrient data to link our samples to water masses and estimate the average age of sampled water in years since equilibration with the atmosphere using CFC-11, CFC-12, SF_6_ (*29*) and DI^14^C concentrations (*30*) for younger and older water respectively.

**Fig. 1.**
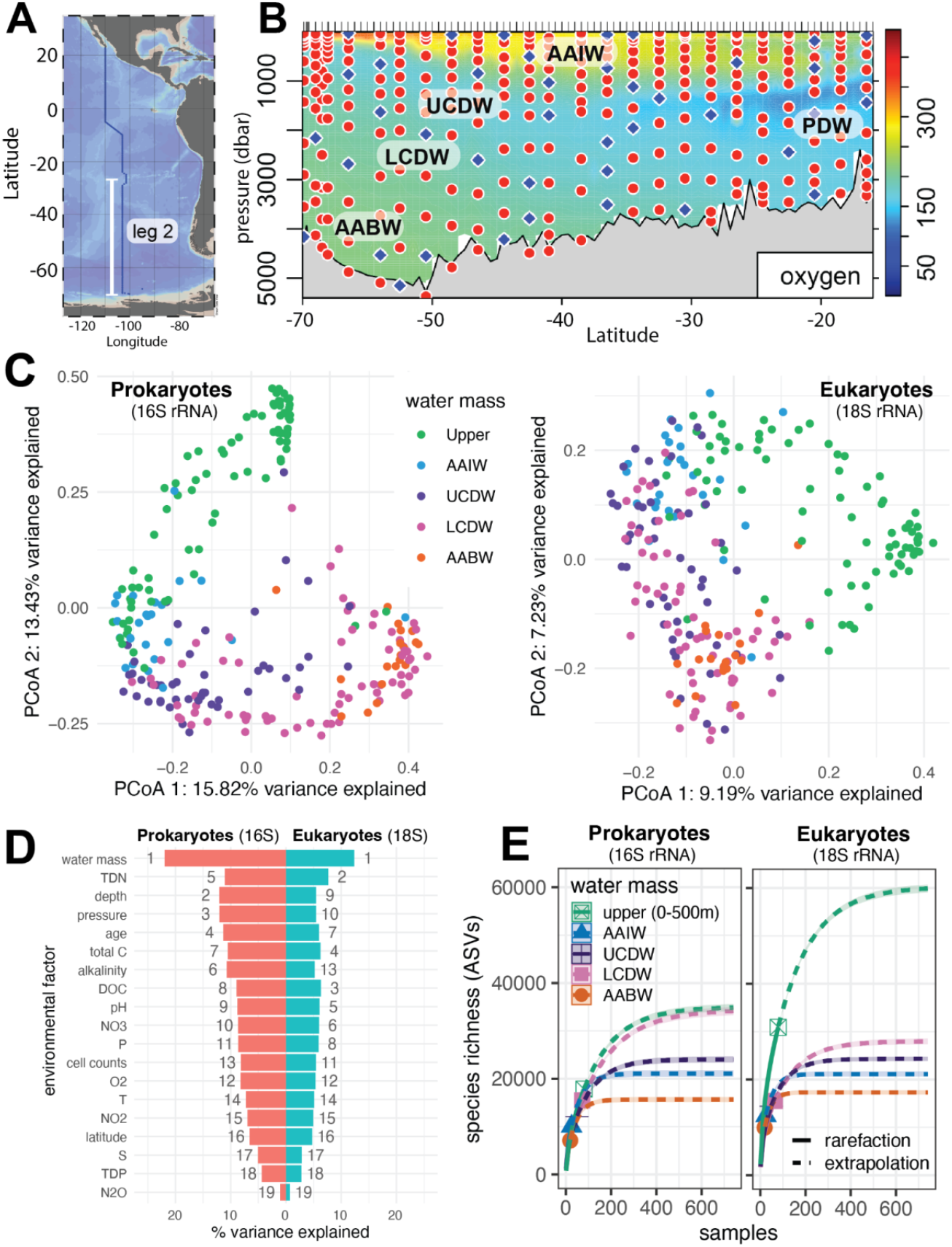
Water mass sampling and microbial biogeography. (**A**) GO-SHIP P18 line with leg 2, in which molecular samples were collected, labeled in white. Generated using Ocean Data View. (**B**) Samples processed for 16S and 18S metabarcoding analysis (red dots) and reserved for metagenomes (blue diamonds) overlay a leg 2 oxygen (μM) section with labeled water masses [Antarctic Bottom Water (AABW), Antarctic Intermediate Water (AAIW), Pacific Deep Water (PDW), Upper Circumpolar Deep Water (UCDW), Lower Circumpolar Deep Water (LCDW)]. (**C**) Principal Coordinate Analysis (PCoA) plot of Bray-Curtis dissimilarity of 16S (left panel) and 18S (right panel) rRNA amplicons. Each point represents a single sample and is colored by water mass: Upper water (< 500m), Antarctic Intermediate Water (AAIW), Upper Circumpolar Deep Water (UCDW), Lower Circumpolar Deep Water (LCDW), Antarctic Bottom Water (AABW) (**D**) Metadata variables found to significantly explain differences in community structure for 16S ASVs (pink) and 18S ASVs (blue; PERMANOVA FDR < 0.05) ordered by percent of variance explained. Bars are numbered to indicate rank of each variable in terms of percent variance explained. (**E**) Rarefied (solid lines) and extrapolated (dashed lines; Chao2 estimator) ASV richness projected to 750 samples for each water mass. We predict to have detected 43-45% of 16S ASVs and 36-37% of 18S ASVs.

### Water sampling and genome recovery

We collected 301 samples across 25 stations during leg 2 of the GO-SHIP P18 line (Fig. 1A) creating an oceanic section of the South Pacific spanning the full water column from roughly 26°S to 69°S. Forty samples spanning a wide range of water age, latitude and depth were chosen to resolve the metabolic potential of the community using shotgun metagenomics (Fig. 1B, blue diamonds). We manually curated 206 medium quality (≥ 50% completeness, < 10% contamination) and 101 high quality (> 90% completeness, < 5% contamination) species-level (95% ANI) prokaryotic metagenome assembled genomes (Data S1). The remaining samples were used to characterize the prokaryotic and eukaryotic community structure with 16S and 18S rRNA metabarcoding analysis (Fig. 1B, red dots), from which we recovered 51,285 16S rRNA amplicon sequence variants (ASVs; Data S2) and 175,993 18S rRNA ASVs (Data S3). Additionally, we spiked in known quantities of *Thermus thermophilus* and *Schizosaccharomyces pombe* gDNA as internal standards in order to estimate ASV abundance in terms of rRNA copies per mL of seawater filtered. This allowed us to approximate the absolute spatial profile of a given taxa rather than its relative abundance (*31*).

### Water masses structure pelagic microbial communities

To identify the strongest environmental features (Data S4) shaping communities, we performed a principal coordinate analysis (PcoA) for both prokaryotic (Fig. 1C, left) and eukaryotic (Fig. 1C, right) amplicons. In both cases, water mass explains more variance in community structure than any other variable measured (Fig. 1D, Data S5), in contrast with the current global paradigm of temperature being the most important driver of community structure in the mesopelagic and above (*8*). This may be because we were able to extensively sample below the mixed layer, where microbes do not encounter large temperature variations. Indeed, 16S rRNA surveys reaching the bathypelagic in the Southern Ocean (*32*) and North Atlantic also found water mass-specific clustering of bacterial communities (*33*). In addition to harboring distinct microbial populations, we also observe that water masses have differing functional capacities (Supplementary text). Other factors that have historically been shown to structure prokaryotic communities in the mesopelagic and above, such as oxygen (*34*), macronutrients (*35, 36*), and latitude (*37*), still hold explanatory power here, but explain less community variance than water mass of origin, depth, and water age (Fig. 1D). While water masses have a high level of endemism for prokaryotes (Fig. S1-S2), many eukaryotic ASVs (14.7%) are present in every water mass (Fig. S1). A notable exception is upper water (< 500 m), which contains 6,919 endemic ASVs, representing a diversity of phytoplankton that are not present in deeper water (Fig. 1E; Data S6-S7). Similarly, within the euphotic zone, a latitudinal gradient in nitrogen availability is the 2nd most important factor explaining variance in eukaryotic community structure (Fig. 1D, Fig. S3).

### Local species richness is driven by depth and age

When considering water masses as a whole, the overall taxonomic repertoire is greatest in upper water for both prokaryotes and eukaryotes (Fig. 1E). However, this is not the case at the level of individual samples. When considering the number of prokaryotic ASVs in each sample, surface water samples harbor the least diversity, and ASV richness drastically increases descending from the surface through the pycnocline. We call this the “phylocline” (Fig. 2A). Just as the pycnocline represents a rapid increase in water density with depth, the phylocline represents a rapid increase in richness with depth. Strikingly, as with the pycnocline, richness is largely stable below the phylocline to full ocean depths (Fig. S4), making it an important delineation between surface waters and deep waters insulated from mixing. A similar depth-integrated trend in richness has been reported off the California coast (*38*) and in 13 locations across the tropics and subtropics (*39*). This, together with the Tara Oceans report of within-sample richness increasing between the surface and the mesopelagic on a global scale (*8*), indicates that the phylocline is a pervasive feature of pelagic microbial ecology.

**Fig. 2.**
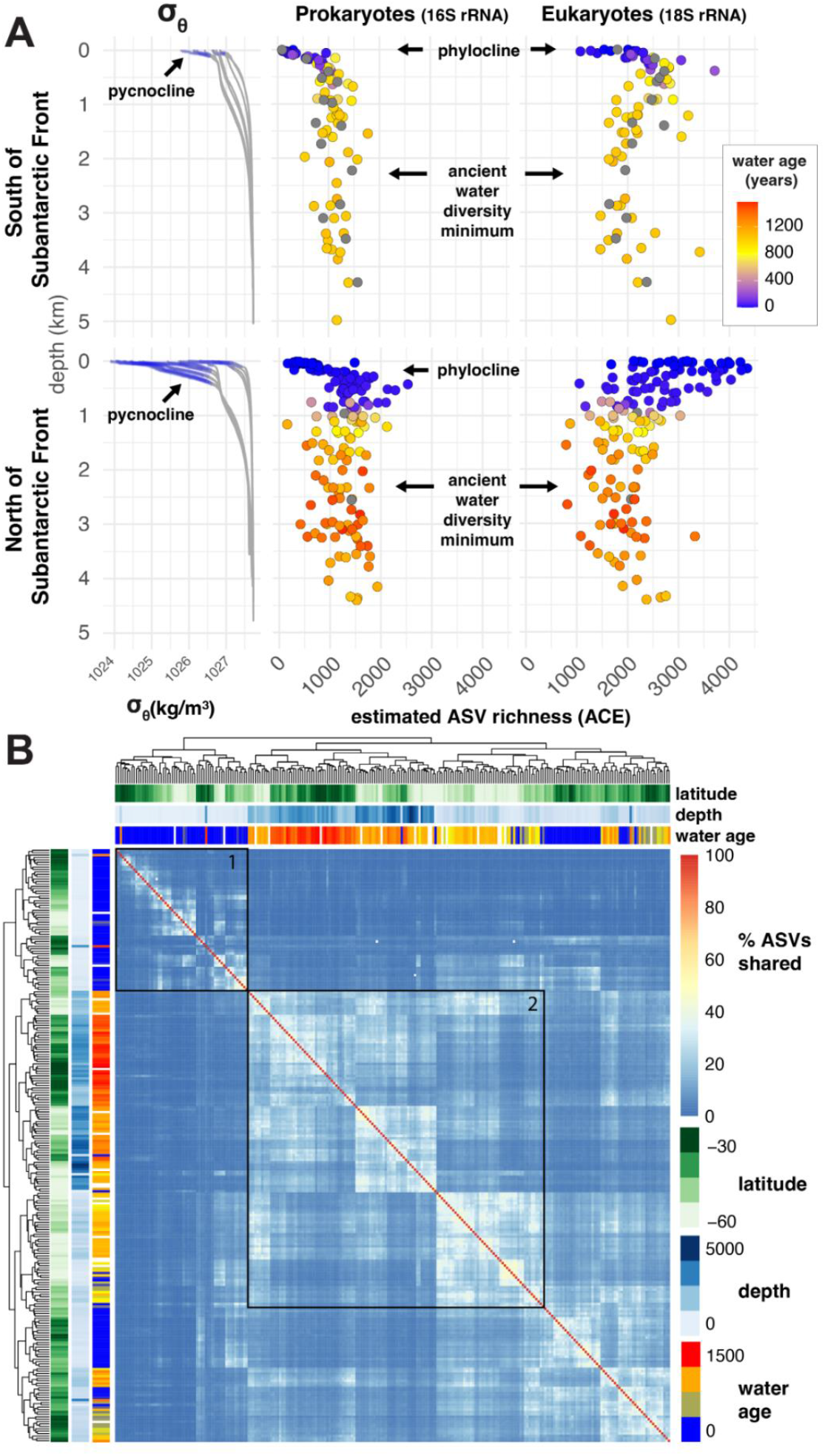
Drivers of species richness. (**A**) Comparison of in-sample diversity across depth to pycnocline. Left: potential density profiles for stations where molecular data was collected. Each station’s pycnocline (defined as 0.01 kg/m^3^ per meter) is highlighted in blue. Middle and right: ASV richness of individual samples (Abundance-based Coverage Estimator; ACE) for prokaryotes (16S; middle) and microeukaryotes (18S; right) colored by water age (samples for which age could not be estimated are in gray). (**B**) Heatmap showing the percent of ASVs (both 16S and 18S) shared between any two given samples, as indicated by the blue to red color bar (right). Samples are hierarchically clustered based on patterns of shared ASVs. Color bars at the top and left of the figure denote the latitude (green) depth in meters (blue) and water age in average years since atmospheric equilibration (orange) of each sample. Samples for which the age could not be estimated are colored white. Box 1 highlights surface samples that have a low percentage of shared ASVs. Box 2 highlights older water samples that have a high percent of shared ASVs.

Lower local species richness at the surface may be explained by regional adaptation to the patchiness of surface water, which experiences fluctuations in temperature and nutrients from which deeper waters are insulated. To test this hypothesis, we compared the percent of ASVs shared between any two given samples (Fig. 2B, Data S8). As predicted, samples clustered into groups of young and old water with young (surface) samples (e.g. Fig. 2B, box 1) sharing fewer ASVs than old samples (e.g. Fig. 2B, box 2). The greater similarity in old water samples explains the apparent dip in prokaryotic richness between 2-4 km, where older water (red) intrudes in our transect. This ancient water minimum is more apparent North of the Subantarctic Front (SAF) of the Antarctic Circumpolar Current (ACC), which contains a higher mixing fraction of older Pacific Deep Water, and is much more dramatic for eukaryotes (Fig. 2A). Prokaryotes may be less impacted by water age than eukaryotes because of their diverse metabolic strategies for thriving in dark, anoxic environments with only recalcitrant carbon sources (e.g. chemoautotrophy, dissimilatory nitrate and sulfate reduction, methane metabolism (*11*))

Microbial eukaryotic richness also forms a phylocline, but only South of the SAF (Fig. 2A). North of the SAF, eukaryotic diversity has the opposite trend: high at the surface and gradually decreasing. This inversion may be explained by the nutrient regimes set up by the ACC. South of the SAF, the ACC drives upwelling that supports large phytoplankton blooms, in which a few species dominate (*40*). The biomass generated by these blooms appears to support a larger diversity of microbial eukaryotes in the mesopelagic (Fig. 2A). North of the SAF, however, downwelling limits nutrients (*41*), driving competition and leading to relatively high surface diversity that gradually declines with depth.

### Microbial genomes self-assemble into spatially-distinct cohorts

To investigate the spatial bounds of individual species, we used Weighted Gene Correlation Network Analysis (WGCNA) to group prokaryotic genomes present in at least 3 samples based only on the similarity of their abundance patterns over latitude and depth. Six biogeographical cohorts emerged (Fig. 3A). Strikingly, microbial cohorts were geographically segregated into regions that can be explained by the prevailing oceanographic and biogeochemical features of the transect (Fig. 3B): a Surface Cohort (pink), dominated by Gammaproteobacteria and Alphaproteobacteria; a Mesopelagic Cohort (orange) dominated by Nitrososphaeria and Gammaproteobacteria genomes; an Antarctic Bottom Water (AABW) Cohort (teal), which follows the sinking of AABW, dominated by Acidimicrobiia and Poseidoniia genomes; an Upper Circumpolar Deep Water (UCDW) Cohort (yellow), located in the mixing zone of North Atlantic Deep Water (NADW) and the Antarctic Circumpolar Current (ACC) and dominated by Gammaproteobacteria and Acidimicrobiia genomes; an Ancient Water Cohort (blue), which is most abundant in water >1000 years old and dominated by Dehaloccoidiia and SAR324; and a Deep Water Cohort (violet), which is abundant below 2 km across all latitudes and is dominated by Nitrososphaeria and Alphaproteobacteria genomes. Clustering 16S and 18S ASVs recapitulated largely the same biogeographical provinces as prokaryotic genomes, but the inclusion of eukaryotes and increased spatial resolution due to larger sample numbers revealed more nuance in the upper ocean (Supplemental text, Fig. S5-S7, Data S2-S3). We also created an interactive tool, the Microbial Ocean Atlas for Niche Analysis (MOANA), that allows users to visualize the distribution of any individual ASV, taxonomic group, or genome across latitude and depth (Supplemental text, Fig. S8). MOANA plots microbial abundance over an interpolated section of any of the environmental metadata collected on the GO-SHIP P18 line, allowing researchers to better understand how their species of interest responds to the overarching physical and nutrient regimes of the South Pacific.

**Fig. 3.**
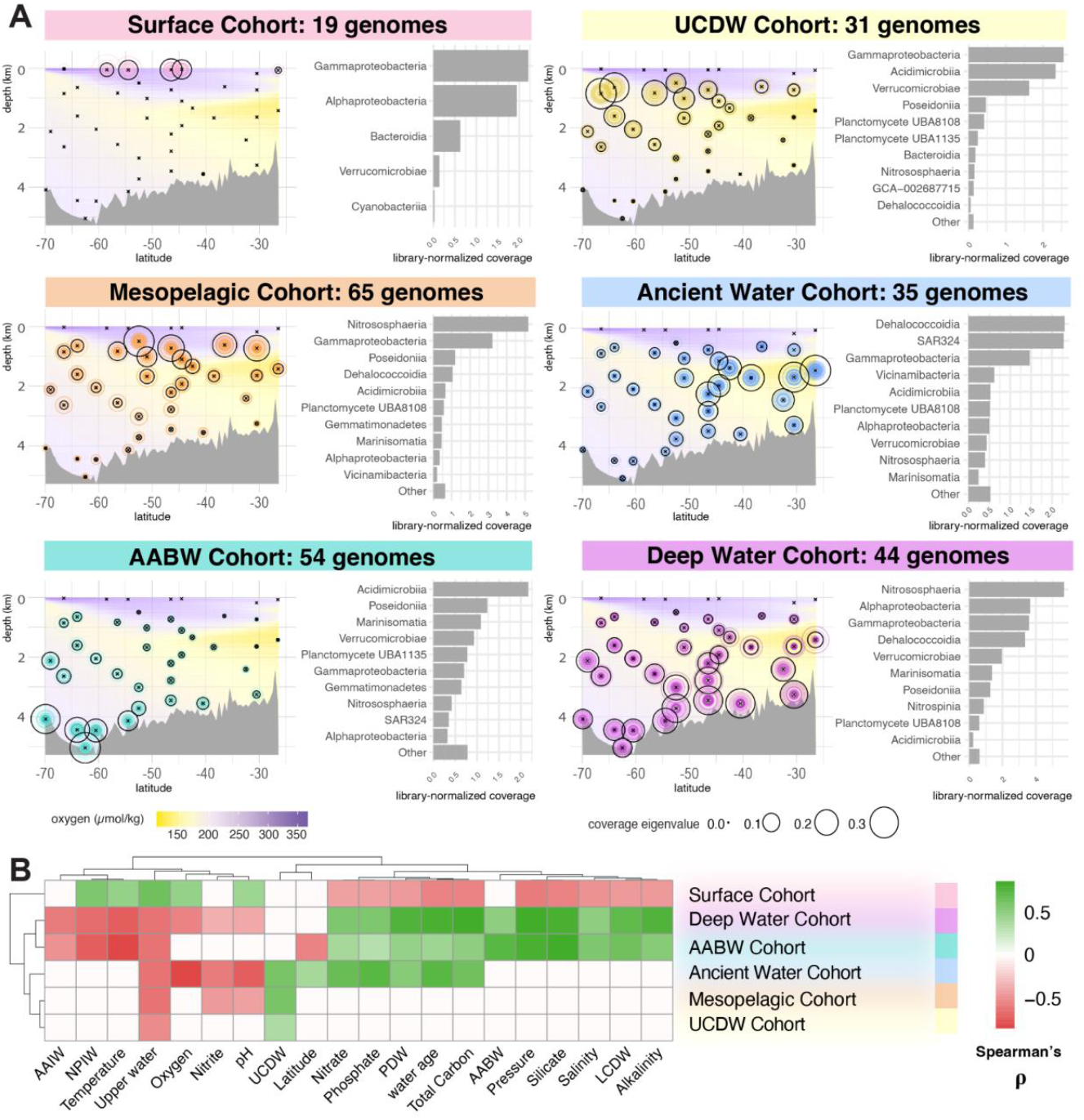
Cohorts of co-occurring microbial genomes as resolved through weighted group correlation network analysis (WGCNA). (**A**) Section plots indicating the location of each microbial province across latitude and depth. Black circles indicate the average abundance profile for genomes in the province and colored circles indicate the abundance of each genome. To the right of each section plot is a bar graph indicating the top 10 taxonomic classes present in the province, by overall library-normalized coverage. (**B**) Heatmap of metadata variables found to significantly correlate with the location of each province (FDR < 0.05) colored by correlation coefficient (Spearman’s rho). Variables with no significant correlation are left blank.

### Genome functional potential is partitioned by oceanographic features

To investigate the functional biogeography of the South Pacific, we repeated WGCNA on KEGG Orthologies (KOs) annotated from genomes, revealing 10 functional zones (Fig. 4, Fig. S9, Data S9). Functional zones are broadly co-localized with genome cohorts, but include additional localities (Supplementary Text). For example, there are six functional zones at the surface that are bounded by wind-driven circulation regimes (e.g. the Subantarctic Front, Polar Front, Southern Antarctic Circumpolar Current Front, and the Oligotrophic subtropical gyre), which establish regions of upwelling and downwelling. Functions broadly important for survival in the surface ocean, such as rhodopsin, carotenoid metabolism, iron acquisition, and heme biosynthesis are ubiquitous at the surface across latitudes (Fig. 4A, Zone 5, light red).

**Fig. 4.**
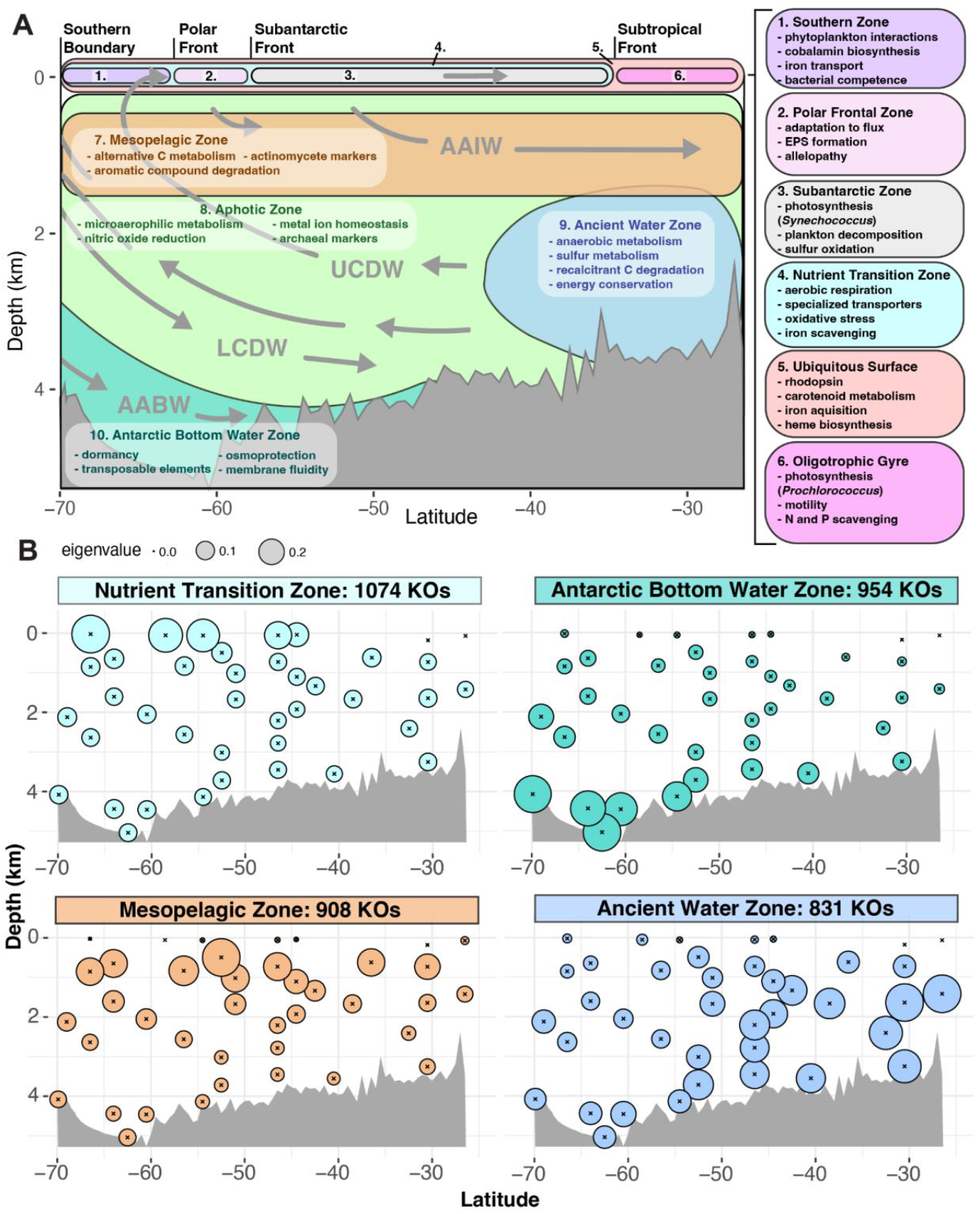
Microbial functional zones of the South Pacific. (**A**) Cartoon summary of zones of co-localized microbial gene functions as resolved through weighted group correlation network analysis (WGCNA) of KEGG Orthologies (KOs) from curated genomes. (**B**) Section plots indicating average coverage pattern (eigensections) of the four functional zones encompassing the most KOs. Section plots for all other functional zones can be found in Fig. S9.

The zones capturing the most functional diversity are the Nutrient Transition Zone (1,074 KOs), Antarctic Bottom Water Zone (954 KOs), Mesopelagic Zone (908 KOs) and Ancient Water Zone (831 KOs; Fig 4B). Functions in the Nutrient Transition Zone (Fig. 4, Zone 4, turquoise) are most abundant in surface waters around Antarctica, and decrease moving northward. They include markers of aerobic respiration, specialized transporters, iron scavenging genes, and oxidative stress response genes. Functions in the Mesopelagic Zone (Fig. 4, Zone 7, orange) are most abundant in the mesopelagic and upper bathypelagic and are associated with degradation of alternative C sources (e.g. 2,5-Furandicarboxylate decarboxylase (*42*), 2-aminomuconate deaminase (*43*), benzil reductase (*44*)) and aromatic compounds.

Functions in the Antarctic Bottom Water Zone (Fig. 4, Zone 10, teal) are most abundant in regions where high-salinity AABW forms and sinks around Antarctica (*45*). They include osmoprotective genes (e.g. *ompR* (*46*)) as well as genes to maintain membrane fluidity across high pressure and cold temperatures. Transposases are also about three times more abundant in AABW (Fig. S10) indicating that surviving cells may horizontally acquire the functions needed to adapt to rapid downwelling. Additionally, several of the bacterial toxin/antitoxin systems in this zone that are typically considered mobile genetic element (MGE) maintenance machinery have been shown to trigger reversible bacteriostasis under stress conditions such as starvation (*47*) and temperature shock (*48*). Such dormancy could confer an adaptive advantage for cells sinking as part of AABW formation. This is not unprecedented, as psychrophilic and psychrotolerant bacteria have been shown to use plasmids to rapidly adapt to cold (*49*). Transposases are also enriched in phytoplankton scaffolds found in AABW (K07497, FDR = 4.39e-5, Data S10), possibly because algae adapted to survive below the photic zone have genomes large enough to tolerate viral ballast.

Functions in the Ancient Water Zone (Fig. 4, Zone 9, blue) are most abundant in the oxygen minimum associated with Pacific Deep Water, the oldest water on earth (*50*). Coverage differences between the origin and terminus of replication (index of replication; iRep) (*51*) estimate that even in water older than 1000 years, approximately 79% of genomes are replicating (Data S11), although more slowly than in surface water (Tukey’s *q* = 0.018; Fig. S11). Functions in this zone are associated with anaerobic metabolism, energy conservation, and the degradation of recalcitrant carbon sources, such as phenolic compounds (*52*).

### Conclusions and Future Directions

Incorporating molecular measurements into the GO-SHIP P18 line allowed us to observe how circulation structures microbial communities. Broadly, we observe a “phylocline” -- a steep transition from low local species richness at the surface to consistently high richness below the mixed layer. We found that both prokaryotic and eukaryotic communities are most easily distinguishable by the water mass they are sampled from. Water age is also important and creates a diversity minimum between 2 and 4 km deep. Individual taxa often have broad, contiguous habitats rather than patchy distributions. While each species has a unique abundance profile, 90% of genomes present in at least 3 samples (n= 225) self-organize into one of six spatial cohorts bounded by depth (Surface, Mesopelagic and Deep Cohorts), large-scale mixing (Upper Circumpolar Deep Water and Antarctic Bottom Water Cohorts), and water age (Ancient Water Cohort). The functional potential of genomes is partitioned by convection, depth, and water age below the mixed layer as well as by wind-driven circulation at the surface. This analysis represents an ocean-basin scale delineation of microbial ecosystems driven by genomic potential and informed by high-resolution oceanographic metadata. We create a baseline of how microbes are structured by GOC, which may slow with climate change (*53*). Collectively, these results illustrate how including depth-integrated molecular measurements in hydrographic sampling programs could create a global, species-resolved atlas of biodiversity and offer biogeochemical insights into the ecosystem services provided by pelagic microbes.

## Acknowledgments

We thank the GO-SHIP program, especially L. D. Talley and P18 Chief Scientists B. R. Carter and R. E. Sonnerup, for generously accommodating ancillary programs. We also thank the crew of the Ronald H. Brown for their commitment to enabling scientific research and C. A. Garcia for onboard sample processing consultations. We thank J. J. Minich for his guidance in the processing of low-yield deep sea samples, D. Kaul and R. H. Lampe for bioinformatic consultation, J. A. F. Giammona for guidance in deploying the Microbial Ocean Atlas for Niche Analysis (MOANA), and K. E. Selph for processing flow cytometry samples. We also thank F. Azam, D. Bartlett, P. Franks and K. Zengler for thoughtful feedback and discussions during the development of this research.

## Funding

US NSF GO-SHIP grant NSF 2023545 (SGP, RES)

Simons Foundation Collaboration on Principles of Microbial Ecosystems (PriME) Grant 970820 (AEA)

NSF graduate research fellowship DGE-1144086 (BK)

NIH Shared Instrumentation Grant 1S10OD010786-01 (DNA Technologies and Expression Analysis Cores at the UC Davis Genome Center)

NIH S10 OD018174 Instrumentation Grant (QB3 Genomics Sequencing Laboratory) Support from the Emerson Collective and the Chan Zuckerberg Initiative (JFB)

## Author contributions

Conceptualization: BCK, RS, EEA, JFB, AEA

Methodology: BCK, RS, HZ, ZF, SGP, RES

Investigation: BCK, RS, ZF, ET, SGP, RES, EEA, JFB, AEA

Visualization: BCK

Funding acquisition: SGP, RES, EEA, JFB, AEA

Writing: The manuscript was primarily written by BCK, with substantial input from JFB and AEA. All authors contributed to the final manuscript.

## Competing interests

JFB is a cofounder of Metagenomi. All other authors declare that they have no competing interests.

## Data and materials availability

All genomes are available on ggkbase at ggkbase.berkeley.edu/GOSHIP-P18-pelagic-marine-genomes. Genomes that are >90% complete have been deposited to the National Center for Biotechnology Information (NCBI) along with shotgun metagenomic and amplicon reads and will be released upon publication. Physical and chemical data collected on the P18 line can be accessed at the CLIVAR and Carbon Hydrographic Data Office (CCHDO) website (https://cchdo.ucsd.edu/cruise/33RO20161119). All other data are available in the main text or supplementary materials.

## Supplementary Materials

Materials and Methods

Supplementary Text

Figs. S1 to S16

Table S1 References (*54-153*)

Data S1 to S16

## Notes

https://ggkbase.berkeley.edu/GOSHIP-P18-pelagic-marine-genomes/organisms

https://hub.docker.com/r/bkolody/moana-p18-prokaryotes

https://hub.docker.com/r/bkolody/moana-p18-eukaryotes

https://hub.docker.com/r/bkolody/moana-p18-genomes

