## Supplemental Text for "Overturning circulation structures the microbial functional landscape of the South Pacific"

**The PDF file includes:**

Materials and Methods
Supplementary Text
Figs. S1 to S16
Table S1 to S6
References 54 - 155

**Other Supplementary Materials for this manuscript include the following:**

Data S1 to S9

### Materials and Methods

Sample collection

Sampling was conducted on board the NOAA Ship *Ronald H. Brown* from January 2-29, 2017 during the second leg of the Global Ocean Ship-Based Hydrographic Investigations Program (GO-SHIP) P18 line. This cruise followed 103$^{o}$W from 26$^{o}$ 29.995'S to 69$^{o}$ 0.014'S, sampling approximately every half degree of latitude. Each deployment consisted of lowering a Sea-Bird Electronics SBE9plus CTD, connected to a 24-place SBE32 carousel, to within 8-12 meters of the bottom using the altimeter on the CTD-rosette package (with the exception of 3 stations). Molecular samples were collected approximately every 4$^{th}$ station (~2 degrees of latitude) at a full range of water column depths. A total of 301 samples were taken over 25 stations. At each station, ~12 sampling depths were chosen to target the major water mass features predicted by the previous P18 iteration, incorporate as wide a range as possible of pressure conditions, and maximize available water.

Preservation of cellular material

Seawater collected for molecular analyses was filtered, over ice, into 5 um and 0.22 um size fractions to capture particle associated and free-living microbes, respectively. Results presented here consider both size fractions together. Volumes filtered ranged from 3-8 liters depending on availability. Filters were immediately submerged in RNAlater$^{TM}$ and stored in liquid nitrogen. In addition, 1 mL of seawater from each sample was preserved in 1% (final concentration) paraformaldehyde for flow cytometry. All samples were preserved in liquid nitrogen throughout transit before being transferred to a -80$^{o}$C freezer. Between samples, the filtration rig and collection jugs were triple-rinsed with 10% bleach, followed by 0.22 um sterile filtered water and sample water.

Library Preparation and Sequencing

For molecular barcoding analyses, half of each filter was used for RNA extraction using the NucleoMag RNA kit (Macherey-Nagel) with an epMotion TMX automated liquid handling system (Eppendorf). *Schizosaccharomyces pombe* gDNA (ATCC 356 #24843D-5, Manassas, VA, USA) and *Thermus thermophilus* gDNA (ATCC 27634D-5) with known rRNA gene copy numbers were used as internal standards for 18S and 16S analyses, respectively (Lin et al., 2018). Internal standards were spiked in after the cDNA synthesis. Samples were grouped into 3 categories based on collection pressure (0-200 dbar, 200-500 dbar, >500 dbar) as a proxy for biomass in order to allow spike-ins to account for ~1% of sample reads. ZymoBIOMICS whole-cell microbial community standards (D6300) were used as a control and were serially diluted to 10^9, 10^7, 10^5, 10^3, 10^2, 10, 1, and 0 cells/mL with ZymoBIOMICS DNA/RNA Shield™ (Cat. No. R1100-50).

The V4-V5 region of 16S was targeted with the degenerate primer pair F515 (GTGNCAGCMGCCG CGGTAA) and R926 (CCGYCAATTYMTTTRAGTTT)(*54*, *55*), which were also used in the *Tara* global oceans sampling program (*56*). The V9 region of 18S rRNA and rDNA were amplified with the primers 1389F (5′-TTGTACACACCGCCC-3') and 1510R (5′-CCTTCYGCAGGTTCACCTAC-3'), which capture about 150bp (*57*), also utilized by *Tara* oceans (*56*). 250 riboTag libraries were multiplexed per MiSeq run (~20 million reads/run). 16S samples were sequenced across six MiSeq PE300 runs and 18S samples were sequenced across on three HiSeq4000 PE150 runs.

For metagenomes, DNA was extracted from whole filters using the NucleoMag Plant kit (Macherey-Nagel). 20 ng of DNA from each sample was used as input for the KAPA HyperPlus library prep kit (Roche). The resultant libraries had insert sizes averaging > 1kb and underwent a series of size selections to remove larger fragments. In order to acquire more representative data for relative abundance information, additional libraries were prepared with the remaining DNA from each sample (between 5.9 and 130 ng) and the NEBNext Ultra II DNA Library prep kit was used with covaris shearing to attain ~600 bp inserts. Both sets of libraries were sequenced on with Illumina Novaseq 6000 with 2 x 150 bp paired-end reads, assembled and used to create genomes, but only NEBNext Ultra II DNA libraries were used for mapping. Sequencing was performed at the DNA Technologies and Expression Analysis Cores at the UC Davis Genome Center and QB3 Genomics, UC Berkeley, Berkeley, CA, RRID:SCR_022170.

Shipboard controls

To account for potential contamination from the ship, control samples were taken by swabbing the bench, floor, and bulkhead surfaces of the ship's laboratory; filtering the 0.22-um sterile-filtered tap water that was used for cleaning the filtration apparatus between samples; and filtering the ship's tap water directly. Two 16S ASVs that were present in all the shipboard control samples were labeled contamination and removed from further analyses. For metagenomic analyses, shipboard control genomes were dereplicated at 95% identity (dRep, default parameters (*58*)) and were annotated with GTDB-Tk (*59*) as *Schlegelella* sp., Rhodocyclaceae sp., *Erythrobacter cryptus*, *Pseudomonas guguanensis*, *Novosphingobium meiothermophilum*, *Cyclobacteriaceae sp. UBA2336*, *Burkholderiales SG8-41* and *Oleibacter sp002733645*. All dereplicated shipboard control genomes originated from either the shipboard tap water or 0.22 um filtered tap water samples and were dominated by freshwater thermophiles with closest relatives isolated from hot springs, which is consistent with them originating from the ship's hot water tank. All shipboard control genomes were included in the final dereplication of marine genomes, and one marine genome (GOSHIP-P18_133_1_6_39S_2409m_maxbin2_1; *Pseudomonas guguanensis*) clustering with a shipboard control genome was removed. As an additional precaution, all shipboard control reads were mapped to the final set of marine genomes and four bins with control reads mapping across more than 10% of the genome at 95% identity or greater were removed. These genomes were annotated as *Alteromonas mediterranea, Alteromonas naphthalenivorans*, *Halomonas aquamarina*, and *Sulfitobacter pontiacus*.

Amplicon analyses

Ribosomal RNA reads were processed using qiime2 (*60*). Sequences were trimmed to remove primers with cutadapt (*61*), and amplicon sequence variants (ASVs) were called using DADA2 (*62*) with custom forward and reverse read trimming based on quality score visualizations. Taxonomy was assigned using qiime feature classifiers. For the 16S amplicon, a naive bayes feature classifier was trained on SILVA138 (www.arb-silva.de), using the 99% identity clustered core alignment file. For the 18S amplicon, a naive bayes classifier was trained on the protist ribosomal database (*63*). Mitochondrial and chloroplast sequences were filtered out.

Because of the low biomass nature of deep ocean samples, serially diluted mock community controls were used to determine the minimal coverage necessary to accurately capture community composition. The percent of ASVs mapping to the correct mock community genera in each dilution was calculated, and an allosteric sigmoidal model was used to determine the coverage level at which 50% of ASVs map to the mock community ($K_{1/2}$) (*64*). The $K_{1/2}$ was determined to be 530 reads for 16S and 96,300 for 18S. Samples below this coverage threshold were removed from downstream analysis (n= 29 for 16S; n= 35 for 18S). Additionally, the two 16S ASVs that were present in all the shipboard control samples (678b5b9d41e8e92f769b8c3a674c831b (*Pseudomona*s sp.) and a594a2db0ceea408a3931a4465c05420 (*Rhodococcus* sp.)) were labeled contamination and removed from further analyses.

DNA amplicon copies/mL were approximated as previously described (*65*) using the volume of seawater filtered, percent of spike-in retained and genome size and amplicon copy number of spike-in species, and accounting for the filter being cut in half. RNA amplicon copies/mL were similarly estimated, but multiplied by a factor of 7.5 to account for only 4 uL of the 30uL RNA extract being used for cDNA synthesis before adding the spike-in. ASVs mapping to the spike-in species were then removed. ASVs mapping at greater than 99.5% identity and with <= 1 mismatch to mock community sequences (BLASTn) were also removed.

In order to visualize community structure (Fig. 1C, Fig. S3B), a Bray-Curtis dissimilarity matrix was first calculated using the vegdist function from the R vegan package (*66*) for both 16S and 18S ASVs, then used as input for a PcoA, which was conducted using the phyloseq (*67*) ordinate function using the MDS method. Eigenvalues from the PCoA were extracted to calculate the percentage of variance explained by each axis. To identify the underlying metadata variables driving community structure differences (Fig. 1D, Fig. S3C), a PERMANOVA was performed with the adonis2 function to test the significance of metadata variables in explaining variance in community structure. Benjamini-Hochberg correction was employed to control for the false discovery rate. Rarefaction and extrapolation of species richness (Fig. 1E, Fig. 2A) were performed using the R iNEXT package (*68*) with incidence frequency input data and the default estimator. In order to compare the number of species shared between water masses (Fig. S1), unfolded venn diagrams were visualized with the UpsetR package (*69*). 18 samples were randomly selected from each water mass as input to avoid a bias towards water masses that were more deeply sampled, and species only annotated at the domain level were removed.

Weighted Group Correlation Network Analysis (WGCNA; Fig. S5 - S7) was performed using 16S and 18S ASVs present in at least 3 samples (*70*). Cohorts were required to include at least 500 ASVs. A soft power threshold of 5 and a module eigengene dissimilarity threshold of 0.92 were used.

To test the significance of the correlation between depth and estimated ASV richness (Fig. S4) and visualize the resulting Pearson correlation, ggscatter() from the ggpubr (*71*) package was used (version 0.6.0).

<u>Genome-resolved metagenomics</u>

Illumina PE150 reads were processed into annotated scaffolds following the ggkbase metagenomic data preparation pipeline (https://ggkbase-help.berkeley.edu/overview/data-preparation-metagenome/), including adaptor and illumina trace contaminant removal with BBTools (*72*), quality trimming with Sickle (*73*), assembly with idba_ud (*74*), gene prediction with prodigal (*75*), and annotation with usearch against KEGG (*76*), UniRef100 (*77*), and UniProt (*78*) databases. Scaffolds from each NovaSeq run were binned using the ggBin snakemake pipeline, which includes sample cross-mapping with bbmap (*72*), following by binning with CONCOCT (*79*), maxbin2 (*80*), MetaBat2 (*81*), and VAMB (*82*), then choosing the best overall bin set with DAS Tool (*83*). Shipboard control reads were processed using the same pipeline.

A total of 5,051 DAS Tool genomes, including both sample and shipboard control genomes, were used together as input for dRep (*58*), which chose 327 species level (95% ANI) representative genomes based on completeness, contamination and N50. One genome clustering with a shipboard control genome was removed. As an additional precaution, shipboard control reads were mapped to marine genomes with bbmap (*72*) and four genomes that recruited control reads at >=95% ID and >10% minimum covered fraction were removed. The dereplicated marine genome set was further manually inspected using ggkbase's interactive binning tool and curated to remove contigs with anomalous GC and coverage as well as obvious taxonomic contamination. Curated genomes were evaluated with checkM (*84*), and a final set of 206 medium quality draft genomes (>= 50% completeness, < 10% contamination) and 101 high quality draft ( > 90% completeness, < 5% contamination) genomes was retained.

The final genome set was further annotated with metabolic-G (*85*) and DRAM (*86*), with default parameters except a minimum contig size of 1 kb for DRAM. Genome coverage was calculated by mapping sample reads to genomes with bbmap and summarizing with coverm. A genome was considered present in a sample if at least 60% of the genome was covered.

To identify genomes enriched in water masses, a Fisher's exact test was used to compare the number of samples containing a given genome that are present in each water mass versus other water masses. Genomes with FDR < 0.05 were said to be enriched in the water mass tested. A subsequent Fisher's Exact test was used to identify KEGG Orthologies (KOs) enriched within genomes significantly associated with each water mass (Fig. S13, Fig. S14).

Similarly, genomes enriched in water of a given age (< 50 years old, 50-1000 years old, or > 1000 years old) were determined using a Fisher's exact test comparing the number of samples containing a given genome that are present in age category versus other age categories. Genome replication rates (Fig. S11) of enriched genomes were estimated using iRep (*51*) with default parameters. Input sam files were generated by mapping reads to binned contigs from the same sample with bowtie2 (*87*) using the "--reorder" flag.

Weighted Group Correlation Network Analysis (WGCNA) was performed (Fig. 3A) using genomes present in at least 3 samples (*70*). A signed adjacency matrix was used and all genomes were required to have coverage values that were positively correlated with the eigenvalues (average coverage pattern) of the cohort they were assigned to. Cohorts were required to include at least 10 genomes. A soft power threshold of 9 and a module eigengene dissimilarity threshold of 0.6 were used. Pairwise correlations (Fig. 3B) were calculated between each cohort's eigenvalues and metadata variables (e.g. oxygen, nitrate, etc.) and tested for significance using the corr.test() function of the R psych (*88*) package. P-values were FDR adjusted using p.adjust().

To establish functional zones (Fig. 4, Fig. S9), library normalized coverage of genomes containing the KO of interest was summed at each location. KOs detected in at least 3 samples were used for WGCNA, using a soft power threshold of 16 and a signed adjacency matrix, requiring KOs to be positively correlated with their assigned module. Modules (functional zones) were required to have at least 200 KOs.

Number of transposases per genome were tallied based on DRAM functional annotations (Fig. S10). A Kruskal-Wallis test and Dunn's Post-hoc test were used to determine whether the number of transposases detected in each genome is significantly different across water masses. Water mass pertainance is defined here as significant enrichment of a genome in a given water mass (Fisher's Exact test, as described above). FSA (*89*) version 0.9.5 was used to implement statistical tests.

Annotation of eukaryotic scaffolds

Genomic scaffolds annotated as eukaryotic were further classified by BLASTX against the PhyloDB v1.077 (*90*) and assigned a Lineage Probability Index (*91*). Repetitive sequences were masked using Red v2.0.0 (*92*), and then gene models were predicted by Augustus v3.5.0 (*93*) with multiple species settings from a wide range of lineages (arabidopsis, candida_albicans, Chlamydomonas_eustigma, ciona, Cyclotella_cryptica, galdieria, Paramecium_tetraurelia, saccharomyces, zebrafish). Species settings, for which the resulting gene models showed lower completeness or higher fragmentation based on BUSCO v5.7.1(protein mode, eukaryota_odb10; (*94*) , were removed, and then overlapping gene models were collapsed into gene loci, with identifiers stored as gene-locus_isoform. Finally, the resulting set of proteins including isoforms was annotated against KEGG v30-01-2023 using KofamKOALA (*95*). For downstream analyses, only the best scoring annotation was kept for each gene.

Complementary Data Collections

Physical and chemical data collected on the P18 line (*96*) can be accessed at the CLIVAR and Carbon Hydrographic Data Office (CCHDO) website (https://cchdo.ucsd.edu/cruise/33RO20161119). Pressure, temperature, salinity, and oxygen were collected continuously with each cast. The CTD system also incorporated an altimeter, transmissometer, fluorometer/backscatter (FLBB) sensor, a Lowered Acoustic Doppler Profiler (LADCP) for measuring velocity profiles, and Chipods for measuring fine-scale temperature structure.

Bottle data collected on this cruise track includes oxygen and salinity, CFC-11, CFC-12, $SF_6$, silicate, nitrate, nitrite, phosphate, pH, and total alkalinity, collected at 24-depths per cast. $^3$He, Ne, tritium, $N_2O$, stable gasses ($N_2$, $O_2$, Ar), dissolved organic carbon-14 ($DO^{14}C$), dissolved inorganic carbon-14 ($DI^{14}C$), particulate organic carbon (POC), black carbon, and rare earth elements were also measured secondarily. POM samples were collected for POC, particulate organic nitrogen (PON), particulate organic phosphorus (POP) and biological oxygen demand (BOD) from the flow-through system at a total of 198 stations.

Cell Counts

*Prochlorococcus* spp., *Synechococcus* spp., heterotrophic bacteria and photosynthetic eukaryotes were enumerated via flow cytometry at the SOEST Flow Cytometry Facility (www.soest.hawaii.edu/sfcf). Samples were thawed in batches and stained as previously described (*97*, *98*) with 1 µg/ml final concentration Hoechst 34442. The flow cytometer used was a Beckman-Coulter Altra mated to a Harvard Apparatus syringe pump for quantitative analyses, equipped with two argon ion lasers, tuned to UV (200 mW) and 488 nm (1 W) excitation. Scatter (side and forward) and fluorescence signals were collected using filters as appropriate, including those for Hoechst-bound DNA, phycoerythrin and chlorophyll. The data generated was in the form of listmode files (FCS 2.0 format) acquired from the flow cytometer using Expo32 software (Beckman-Coulter). Population designations, based on the scatter and fluorescence signals, were generated from the listmode files using FlowJo software (Tree Star, Inc., www.flowjo.com).

Water mass classification

Optimum MultiParameter (OMP) analysis (*99*, *100*) was used to determine the mixing fractions of the water masses present in each sample. This method uses conservative water mass properties (temperature, salinity) as well as quasi-conservative properties (oxygen and nutrients, taken together as a single variable) to determine the proportion of water in each sample originating from various water masses.

Established temperature, salinity, and nutrient characteristics (*45*) were used to choose precise coordinates and depths/potential densities for water mass end members (Table S5). The remaining water mass characteristics (salinity, potential temperature, oxygen, phosphate, nitrate, and silicate) were then determined at the given location using the World Ocean Circulation Experiment (WOCE) Global Hydrographic Climatology (downloaded 1/15/19, last modified 12/23/2010) data, parsed using the R ncdf4 package (*101*). For Antarctic Bottom Water (AABW), North Pacific Intermediate Water (NPIW), and Antarctic Intermediate Water (AAIW), defining latitudes, longitudes, potential densities and potential vorticities were adapted from (*102*). For upper and Lower Circumpolar Deep Water (LCDW), a high nutrient oxygen minimum layer and a low nutrient salinity maximum layer, respectively (*45*, *103*–*106*), were searched for on WOCE Pacific section plots in order to define water mass coordinates and depths.

Upper ocean circulatory features were defined based on definitions in Orsi et al. (*107*). The latitude of the Polar Front was defined as the northern limit of 2$^{o}$C Antarctic Winter Water at 200 m deep. The Subantarctic Front was defined as 57$^{o}$S at the northernmost limit of the low salinity intermediate layer. The Subtropical Front was defined as 35$^{o}$S, updated from the position given in Orsi et al. (*107*) based on nitrate and salinity transitions of our cruise track (Fig. S12).

Pycnocline determination

Potential density profiles for each station (Fig. 2A) were calculated from CTD data in R, parsed with ncdf4 (*101*). The R gsw package (*108*) was used to calculate sample depth from pressure and latitude and potential density from salinity, temperature, and pressure. The pycnocline was defined as depths at each station where the vertical potential density gradient exceeds 0.01 kg/m$^3$ per meter.

Water mass age determination

First, CFC-11, CFC-12, and $SF_6$ were used to assign a tracer age to samples wherever possible. Empirically derived CFC-11, CFC-12 (*109*), and $SF_6$ (*103*) solubilities were used to determine the atmospheric concentration of each tracer at the time the given water parcel equilibrated with the atmosphere. These concentrations were then compared with the global mean NOAA atmospheric tracer record and interpolated using the R splines function (R Development Core, 2011) in order to determine the average year that the sampled water was in contact with the atmosphere. This value represents the "tracer age," or weighted average of the water sampled, because in truth, the water is an amalgam of water parcels that equilibrated with the atmosphere at different times. The extent to which the true "advective age" aligns with this tracer age depends on the degree of mixing the water parcel has experienced. The advective age can be laboriously calculated using transit-time distribution (TTD) theory, but doing so is outside of the scope of this project. Instead, tracer-aged samples were compared against TTD calculations made for the previous occupation of the P18 line (*110*) as a measure of quality control.

Water masses older than can be predicted with CFC and SF6 data were aged using dissolved inorganic carbon 14 dating (DI14C). High-precision Δ14DIC measurements were processed at Woods Hole Oceanographic Institution's (WHOI) National Ocean Sciences Accelerator Mass Spectrometry lab (NOSAMS) by the McNichol group using standard AMS (accelerator mass spectrometry) protocols (111–113). Δ14DIC was converted into radiocarbon age using the equation: 14C age = -8033*log(1+(Δ14DIC /1000)) (114). Radiocarbon age was converted into calibrated years before 1950 (cal BP) using the standard marine radiocarbon calibration curve, Marine13 (*115*) interpolated via the R splines function (R Development Core, 2011). Cal BP was constructed for the P18 section and interpolated to the locations of genomics samples using bivariate linear interpolation via the R akima package (*116*).

Data manipulation and Visualization

Data manipulation and visualization was aided by the use of various R (*116*) packages: tidyverse (*117*), plyr (*118*), dplyr (*119*), tidyr (*120*), stringr (*121*), reshape2 (*122*), and ggplot2 (*123*). Cruise transect maps and supplemental sections were generated using Ocean Data View (ODV) for Fig. 1A, Fig. S3A, and Fig. S12 (*124*). Metadata sections were created by interpolating data with the R akima package (*116*) for Fig. 3A, Fig. S6, Fig. S7, and Fig. S8. Sections were generated using the R oce package for Fig. 1B and S16. Water mass sections (Fig. S2) were visualized using a custom matlab script. The R packages pheatmap (*125*) and RColorBrewer (*126*) were used for making heatmaps (Fig. 2B, Fig. 3B).

### Supplementary Text

Microbial Ocean Atlas for Niche Analysis (MOANA)

MOANA (Fig. S8) was built as an R shiny app (*127*) and uses the packages tidyverse (*117*), ggplot2 (*123*), hrbrthemes (*128*), shiny (*129*), shinythemes (*130*), shinyjs (*131*), png (*132*), ggpubr (*71*), grid (*133*), and cowplot (*134*). A ready-to-run docker image of the app is available at https://hub.docker.com/r/bkolody/moana-p18-prokaryotes for prokaryotes (16S rRNA amplicons), https://hub.docker.com/r/bkolody/moana-p18-eukaryotes for eukaryotes (18S rRNA amplicons) and https://hub.docker.com/r/bkolody/moana-p18-genomes for reconstructed genomes.

Follow the steps below to run the application:

1) Make sure docker is installed on your machine (https://docs.docker.com/engine/install/).
2) Download the image from Docker Hub by running the following command in your terminal or command prompt:

   `docker pull bkolody/moana-p18-prokaryotes` (for 16S rRNA abundance)

3) Run the docker container:

   `docker run -p 3838:3838 bkolody/moana-p18-prokaryotes`

   This command maps port 3838 on the container to port 3838 on your local machine, ensuring the app is accessible.

4) Access the application:

   Open a web browser and navigate to: `http://0.0.0.0:3838`

   Alternatively, use: `http://localhost:3838`

MOANA should now be running and accessible in your browser. To stop the application, press Ctrl+C in the terminal running the container, or stop it using Docker Desktop or the Docker CLI. For atlases of 18S rRNA and genome abundance, substitute `bkolody/moana-p18-prokaryotes` with `bkolody/moana-p18-eukaryotes` and `bkolody/moana-p18-genomes`, respectively.

<u>Species-level biogeography of microbial eukaryotes</u>

Three amplicon microbial eukaryote cohorts localized to the surface ocean but formed distinct latitudinal subgroups (Fig. S5). The Southern Surface Subgroup (n = 1,350 ASVs) lies between Antarctica and the Polar Front. Primary producers in this group are known to thrive in the nutrient-rich Southern Ocean, such as a diversity of diatoms (Bacillariophyta), especially *Chaetoceros* and *Hemiaulus* spp., and picobiliphytes. The Antarctic Circumpolar Current (ACC) Subgroup (n = 1,690 ASVs) centers on the mid-latitudes of the transect (~45ºS - 60ºS). In this region, diatoms continue to dominate the eukaryotes, but they include more poorly-annotated species and co-exist with cyanobacteria. The Oligotrophic Surface Subgroup (n = 10,036 ASVs) lies in the northernmost portion of the transect (~26ºS - 50ºS) where nutrients are the most limiting. *Prochlorococcus*, and to a lesser extent, *Synechococcus* are the dominant primary producers. Photosynthetic eukaryotes are less abundant and dominated by small algae with high surface area to volume ratios like pelagomonas and picochlorophytes. Instead, abundant eukaryotic ASVs include diverse bacterivorous species like Strombidiida ciliates and the Pseudofungus MAST-1C (*135*).

An additional three amplicon cohorts occur just below the epipelagic and are differentiated by latitude (Fig. S6). The ACC Midwater Cohort (n= 9,772 ASVs) and Oligotrophic Midwater Cohort (n= 2,211 ASVs) occupy the mid- and northern latitudes of the transect, respectively. Cyanobacteria are abundant in both regions, indicating deep chlorophyll maxima, but are especially dominant in the oligotrophic northern region, as corroborated by cell counts (Fig. S16, Data S4). Other microbes in the Oligotrophic Midwater Cohort are known to have streamlined genomes, such as *Nitrosopumilus*, one of the smallest known cells (*136*). This region also harbors a unique diversity of Stramenopiles and three Telonemia groups. The Southern Ocean Midwater Cohort (n = 1,468 ASVs) is dominated by diverse diplonemids, unclassified diatoms, and RAD-C and RAD-B radiolarians.

The remaining three amplicon cohorts directly co-occur with the Ancient Water (n = 1,714 ASVs), AABW (n = 3,412 ASVs), and UCDW (n = 2,767 ASVs; Fig. S7) cohorts established using genomes. Due to the depth of these regions, photosynthetic eukaryotes in these cohorts are vastly outnumbered by heterotrophic lineages, such as diplonemids, labyrinthulids, and ciliates. The Ancient Water Cohort is distinct from other regions, containing a large number of unidentified ASVs, as well as MAST-1D Pseudofungi.

<u>Functional differences across water masses</u>

To identify prokaryotic genomes enriched in water masses, a Fisher's Exact test was used to compare the number of samples for which a given genome was recovered from each water mass versus other water masses. Fifty-three genomes were enriched in upper water (< 500m), 26 in AAIW, 12 in UCDW, 16 in LCDW, and 41 in AABW (FDR <0.05; Data S1). Significant functional enrichments were mostly found in upper water (n= 206) and AABW (n=26) genomes, with only two functions enriched in UCDW and one in AAIW and LCDW (Data S8).

To investigate the  potential for carbohydrate turnover of South Pacific water masses, a Fisher test was used to identify KEGG Orthologies (KOs) and Carbohydrate-Active enZYmes (CAZys) that were significantly enriched in genomes from each water mass (FDR < 0.05, odds ratio (OR) >1, present in at least 5 genomes). Functions enriched in the upper 500 m compared to other water masses related to light harvesting and photoprotection, iron transport, motility, and metabolism (Fig. S13). Light responsive functions include porphyrin/chlorophyll biosynthesis genes (e.g. menaquinone-dependent protoporphyrinogen oxidase, n= 6 genomes including cyanobacteria), sensory rhodopsin (29 genomes; 55%), and carotenoid metabolism genes (e.g. beta-carotene 15,15'-dioxygenase (32 genomes; 60%), 15-cis-phytoene synthase (36 genomes; 68%)). The enrichment of iron complex outer-membrane receptor proteins (25 genomes; 47%) and manganese/iron transport systems (17 genomes; 32%; Data S9) may be due increased iron demand from photosynthesis and cellular respiration (*137*), as well less bioavailable iron (*138*). The enrichment of flagellar genes in organisms of the upper oceans (30 genomes; 57%) may be due to high hydrostatic pressure and low temperature of the deep ocean, which could  impede motility (*139*). Enrichment of amino acid, nucleotide, carbohydrate, and lipid metabolism genes in the upper ocean likely reflects the substrate-rich environment of the photic zone. Indeed, we only found CAZys involved in the breakdown of arabinan, mixed-linkage glucans, xylans, xyloglucan, and chitin (FDR < 0.05, OR 3.3-7.9; Data S9) enriched in the upper ocean.

Of the KOs enriched in AABW (Fig. S14), 17 were related to mobile genetic element acquisition and maintenance (transposases and putative transposases (n=15), DNA transformation protein (n=1), death on curing protein (n=1)) and three were components toxin-antitoxin systems, HicAB (K07339, K18843), VapBC (K18829) known for conferring cell dormancy under stress conditions such as amino acid and glucose starvation (*47*). The ability to become dormant could confer an adaptive advantage for cells experiencing extreme temperature and pressure changes as part of AABW formation. Other functions include the TetR/AcrR family of chemical sensing transcriptional regulators (K16137), which respond to several signals including osmotic stress (*140*), and 2-Aminoethylphosphonate dioxygenase (K21195), which allows organisms to utilize phosphonates as a phosphorus source.

More than 70% of AABW genomes contained enriched mobile genetic element acquisition and maintenance machinery, pointing to the potential for horizontal gene transfer to enable rapid adaptation to the deep ocean. Death on curing proteins (K07341, n = 17) were only detected in genomes enriched in AABW (47.5%) or with no significant enrichment, on average originating from genomes sampled from 4092 m deep. The genomic context of these scaffolds further indicates their horizontal transfer and utility for deep ocean survival (Table S6). They were often found nearby integrases, phage integrases, and transposases. In 11 of 20 cases, death on curing proteins were associated with toxin-antitoxin systems known for dormancy stress responses, and in all but one case, these systems were either directly up or downstream of the MGE stabilizing protein. The most common dormancy system annotation is mazEF superfamily (n=8), which is an. The MazEF toxin-antitoxin system that can trigger a state of reversible bacteriostasis in response to various stressors (*48*) such as starvation, temperature shock, and DNA damage (*141*). Eight proteins were annotated as antitoxins and one was annotated as a MazEF family AbrB-like protein. ArbB is a B. subtilis gene /protein that regulates gene expression during the transition between vegetative growth and the onset of stationary phase and sporulation. Other toxin/antitoxin systems were the Phd/YefM (n= 1) and hipA (n=1) family type II toxin-antitoxin systems, as well as an undescribed addiction module antidote protein (n=1). Death on curing family proteins were also associated with genes involved in stress response (cold-shock protein, LysR, integration host factor subunit beta), fatty acid metabolism (e.g. Clp protease, enoyl-CoA hydratase, LapA family protein), carbon and nutrient acquisition (tripartite tricarboxylate transporter, serine/threonine dehydratase, xylose isomerase, glucose/arabinose dehydrogenase, sulfatases, formate/nitrite transporter, ammonium transporter, cobalamin-independent methionine synthase, tonB iron transporter), and bacterial competence (DNA uptake protein comEA). It has been previously suggested that an increase in MGE with depth is a product of decreased selection pressure with lower cell concentrations (*142*). Here, however, the strong enrichment of MGEs in AABW paired with the association of MGEs with depth-adaptive functions such as dormancy-enabling toxin-antitoxin systems indicates that MGEs are allowing cells to survive the strong selective pressure of bottom water formation.

Functional Zone 1: Southern Zone

To investigate the functional biogeography of the South Pacific, we repeated WGCNA on Kegg Orthologies (KOs) annotated from genomes, revealing 10 functional zones (Fig. 4, Fig. S9, Data S6). Functions in Zone 1 (Fig. 4A, Fig. S9; purple; Data S6) are most abundant in the Southern Zone of the Antarctic Circumpolar Current (ACC), which lies between the Southern Boundary of the Antarctic Circumpolar Current (ACC) and the Southern ACC Front (*45*) (Fig. S12). In this zone, the upwelling of nutrient-rich Lower Circumpolar Deep Water fuels blooms (*143*) of large phytoplankton (Fig. S15), and functions enriched in this region relate to motility (e.g. flagellar biosynthesis protein, FlhF; K02404; n = 32 genomes) and phytoplankton-bacterial interactions, such as bacterial competence (e.g. ComEA, K02237, n=45 genomes; ComEC, K02238, n= 151 genomes), and virulence (e.g. lemA (*144*), K03744, n = 76 genomes). Vitamin B12 and iron often co-limit phytoplankton growth in the coastal Antarctic(*91*), and here we observe an enrichment of proteins involved in Vitamin B12 biosynthesis (e.g. Thiosulfate sulfurtransferase;

K02439, n= 28 genomes), iron transport (e.g. iron(III) transport system ATP-binding protein, K02010, n = 50 genomes; ferrous-iron efflux pump FieF, K13283, n = 34 genomes) and siderophore production (e.g. outer membrane protein SlyB (*145*); K06077, n = 35 genomes).

<u>Functional Zone 2: Polar Frontal Zone</u>

Functions in Zone 2 (Fig. 4A, Fig. S9; pink; Data S6) are most abundant in the Polar Frontal Zone between the Subantarctic and Polar fronts of the ACC (*45*). This region is characterized by seasonally high primary productivity and fluctuating conditions, as strong eddies of the Polar Front move northward causing mixing (*146*). Functions involved with adaptation to flux are abundant here. For example, a diversity of sugar transporters are present along with sugar fermentation stimulation protein A (K06206; n = 103 genomes), which helps to fine-tune the balance between respiration and fermentation depending on the available sugar types and environmental conditions(*147*). Proteins involved in allelopathy, like enediyne biosynthesis protein E4 (K21162; 124 genomes), PmbA (K03592, 79 genomes)(*148*), macrolide efflux protein DHA3 (K08217, n = 70 genomes), and tetracycline resistance protein DHA1 (K08151, n = 47 genomes) are also abundant, as are proteins involved in the synthesis of exopolysaccharides (EPS), like exopolysaccharide production protein ExoY (K16566, n= 14 genomes) and dTDP-4-dehydrorhamnose 3,5-epimerase (K01790; n = 87 genomes).

<u>Functional Zone 3: Subantarctic Zone</u>

Functions in Zone 3 (Fig. 4A, Fig. S9; grey; Data S6) are most abundant in the Subantarctic Zone, which is bounded in the South by the Subantarctic Front of the ACC and in the North by the Subtropical Front (*45*). This region is characterized by moderate macronutrient concentrations (Fig. S12) but sinks the most carbon in the Southern Ocean (*149*, *150*), with primary productivity an order of magnitude higher than the Polar Frontal Zone (*151*). Heterotrophic bacteria are most abundant in this region, as are *Synechococcus* cells (Fig. S16). This region is enriched in proteins involved in cyanobacterial photosynthesis (e.g. PsaB, K02690, n= 4 genomes) as well as the breakdown of phytoplankton biomass. For example, phytol kinase (chlorophyll degradation, K18678, n= 16 genomes), beta-carotene hydroxylase (pigment degradation, K02294, n= 9 genomes), and uronate dehydrogenase (K18981, n = 14 genomes), which breaks down D-glucuronic acid, found in diatom cell walls (*152*) are enriched in the Subantarctic Zone.

<u>Functional Zone 4: Nutrient Transition Zone</u>

Functions in the Nutrient Transition Zone (Fig. 4, light blue; Data S6) are present in the Southern, Polar Frontal and Subantarctic Zone. They are most abundant in the south and decrease moving northward. They include markers of aerobic respiration, such as cytochrome c biogenesis proteins (K02197 - K1200). This region is also rich in specialized transporters like the TRAP-type transport system periplasmic protein (K21395, 168 genomes) and C4-dicarboxylate transporters (e.g. K1169, K03326, K10125) and iron scavenging genes like iron(III) transporters (K02012, 118 genomes and  K02011, 121 genomes) and putative iron-regulated proteins (e.g. K07231, 21 genomes).  Oxidative stress response genes like DNA repair proteins RadA/Sms (K04485; n= 173 genomes) and formamidopyrimidine-DNA glycosylase (K10563, n= 185 genomes) are also abundant in this zone.

<u>Functional Zone 5: Ubiquitous Surface Zone</u>

Some functions are uniformly enriched across the surface ocean (Fig. 4A, Fig. S9, light red; Data S6). Several are essential genes, such as ribosomal protein S3 (K02982, n=282 genomes). Others support light harvesting, such as those involved in iron acquisition (e.g. heme iron utilization protein K07226, n= 51 genomes; iron complex outer membrane receptor protein K02014, n= 157), carotenoid metabolism (e.g. beta-carotene 15,15'-dioxygenase, K21817, n = 39 genomes) and heme biosynthesis (e.g. uroporphyrinogen decarboxylase, K01599, n = 191 genomes). Sensory rhodopsin is also broadly present in the surface ocean (K04643, n= 39 genomes).

Functional Zone 6: Oligotrophic Gyre Zone

Zone 6 corresponds to the oligotrophic gyre in the Northern edge of the P18 transect (Fig. 4A, Fig. S9, magenta; Data S6). Functions in this region relate to adaptation to nutrient stress and utilization of alternate carbon sources. For example, phosphate and phosphonate transporters (e.g. Phosphate-selective porin OprO and OprP, K07221, n = 96 genomes; Phosphonate transport system permease protein, K02042, n = 72 genomes) are enriched in this region, as well as urease (K01429, n = 45 genomes) and its accessory proteins. Proteins involved in the utilization of alternate carbon sources include xylose isomerase (K01805, n = 84 genomes), gluconolactonase (K01053, n = 135 genomes), UDP-glucuronate decarboxylase (K08678, n = 54 genomes) and hypoxanthine phosphoribosyltransferase (purine salvage. K00760, n=148 genomes). Prochlorococcus is also abundant in this region (cell counts; Fig. S14, Data S4) because of its streamlined genome and low nutrient requirements (*153*).

Functional Zone 7: Mesopelagic Zone

Functions in Zone 7 are most abundant in the Mesopelagic (Fig. 4, orange; Data S6). Notable proteins are involved in the degradation of alternative C sources, aromatic carbon biosynthesis and degradation, and actinomycete markers. For example, scyllo-inositol 2-dehydrogenase (K16044, n = 12 genomes), which degrades inositol, is abundant in this zone, as are enzymes that degrade cyclic ketones (e.g. cyclohexanone monooxygenase, K03379, n= 37 genomes). Enzymes from the Shikimate pathway (e.g. aromatic aminotransferase, K00837, n= 12 genomes), which allows for the biosynthesis of aromatic amino acids, are enriched in this region, possibly allowing cells to produce amino acids that would otherwise be difficult to access in the deep ocean. Proteins involved in the metabolism of aromatics are also enriched in this region (e.g. 2,5-Furandicarboxylate decarboxylase, K16874, n = 12 genomes; 2-iminobutanoate/2-iminopropanoate deaminase and 2-aminomuconate deaminase, K09022,K15067, n= 52 genomes; benzil reductase, K16216, n=13 genomes), possibly allowing cells to degrade less favorable phytoplankton biomass. Actinobacterial-specific markers such as Meromycolic acid enoyl-[acyl-carrier-protein] reductase (K11611, n = 13 genomes) and genes involved the mycothiol oxidative stress response system ( e.g. mycoredoxin, K18917, n= 27 genomes; Mycothiol S-conjugate amidase,K18455, n= 13 genomes) are enriched in this region, and several actinobacterial genomes are exclusively present here (e.g. GOSHIP-P18_129_2_8_29S_1643m_Actinobacteria_66_14, GOSHIP-P18_161_1_14_128S_727m_Actinobacteria_68_26).

Functional Zone 8: Aphotic Zone

Functions in Zone 8 are uniformly abundant below the surface ocean (Fig. 4A, Fig. S9, green; Data S6), and are involved in microaerophilic metabolism (e.g. glycolate oxidase, K00104, n= 150 genomes; 5,10-Methylenetetrahydromethanopterin reductase, K00320, n= 75 genomes; Heterodisulfide reductase, K08264, n= 73 genomes) metal ion homeostasis (e.g. DtxR family Mn -dependent transcriptional regulator, K03709, n = 136 genomes; P-type Cu2+ transporter; K01533,K17686, n= 65 genomes) and nitric oxide reduction (e.g. nitric oxide reductase NorQ, K04748, n=57 genomes). Functions exclusive to archaea are also enriched in this zone, such as archaeosine synthase (K06936, n= 28 genomes) and archaea-specific DNA-binding protein (K03622, n= 14 genomes).

Functional Zone 9: Ancient Water Zone

The Ancient Water Zone (Fig. 4, dark blue; Data S6) corresponds to the oxygen minimum in the Northern edge of the P18 transect associated with Pacific Deep Water, the oldest water on earth. Functions in this zone are associated with anaerobic metabolism (e.g. anaerobic carbon-monoxide dehydrogenase, K00196, n = 9 genomes), energy conservation (e.g. trehalose synthase, K13057, n =17 genomes), and the degradation of recalcitrant carbon. For example, arylsulfate sulfotransferase (K01023, n = 70 genomes) sulfates phenolic compounds and benzylsuccinate CoA-transferase (K07544, n = 37 genomes) allows for the anaerobic degradation of toluene.

Functional Zone 10: Antarctic Bottom Water (AABW) Zone

The AABW Zone (Fig. 4, turquoise; Data S6) is located in the sinking shelf water around Antarctica. It contains osmoprotective genes such as the osmotic stress regulator OmpR (K02483, K07660, n = 12 genomes) and osmoprotectant transport systems (e.g. K05847; n = 12 genomes) that may protect cells from the high salinity of the brine rejection process that forms Antarctic Bottom Water. Genes to maintain membrane fluidity across high pressure and cold temperatures are also most abundant in this region, such as 1-acyl-sn-glycerol-3-phosphate acyltransferase (K00655,K05939; n = 30 genomes). Transposases (e.g. K07491; n = 54 genomes; K07498, n = 25 genomes) are also about three times more abundant in AABW (Fig. S10) indicating that surviving cells horizontally acquire the functions needed to adapt to rapid downwelling. Additionally, there are several toxin/antitoxin systems that are enriched in this zone, such as HicB (K18843, n= 37 genomes), ParE1 (K19092, n = 32 genomes) and HigA (K21498, n = 30 genomes). These may trigger reversible bacteriostasis, conferring an adaptive advantage for cells sinking as part of AABW formation. Sporulation genes like spore maturation protein CgeB (K06320, n = 16 genomes), which is implicated in crust glycosylation (*154*), and the spore coat dehydration genes (*155*) *spmA* and *spmB* (K06373, K06374; n = 10 genomes) are also enriched in this region. These genes are often contained within genomes from poorly described lineages not known for spore formation (e.g. Alphaproteobacteria PTKB01 sp002937555) and could be involved in novel mechanisms for membrane adaptation to high pressure and low temperature.

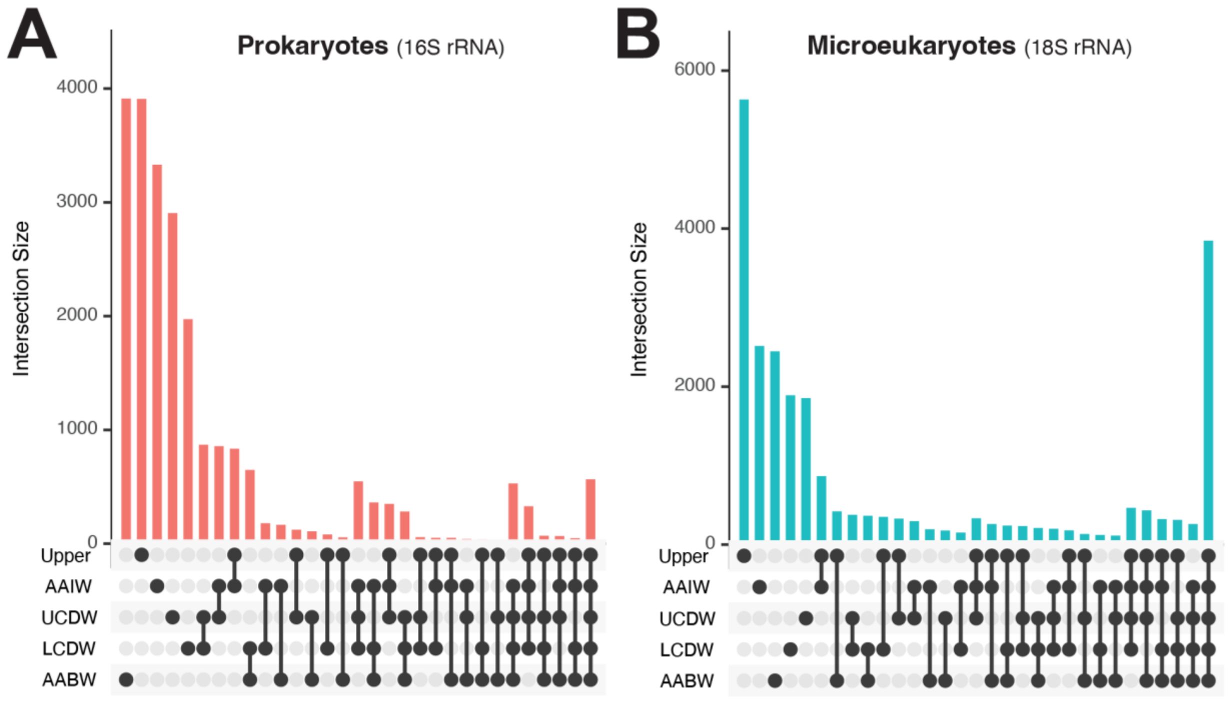


**Fig. S1. Unique and shared ASVs across water masses.**
(A) Upset plot (unfolded venn diagram) showing the number of unique prokaryotic (16S) ASVs shared by each combination of water masses. (B) Upset plot (unfolded venn diagram) showing the number of unique prokaryotic (18S) ASVs shared by each combination of water masses. Eighteen samples were randomly selected to be included  from each water mass in order to avoid oversampling bias.

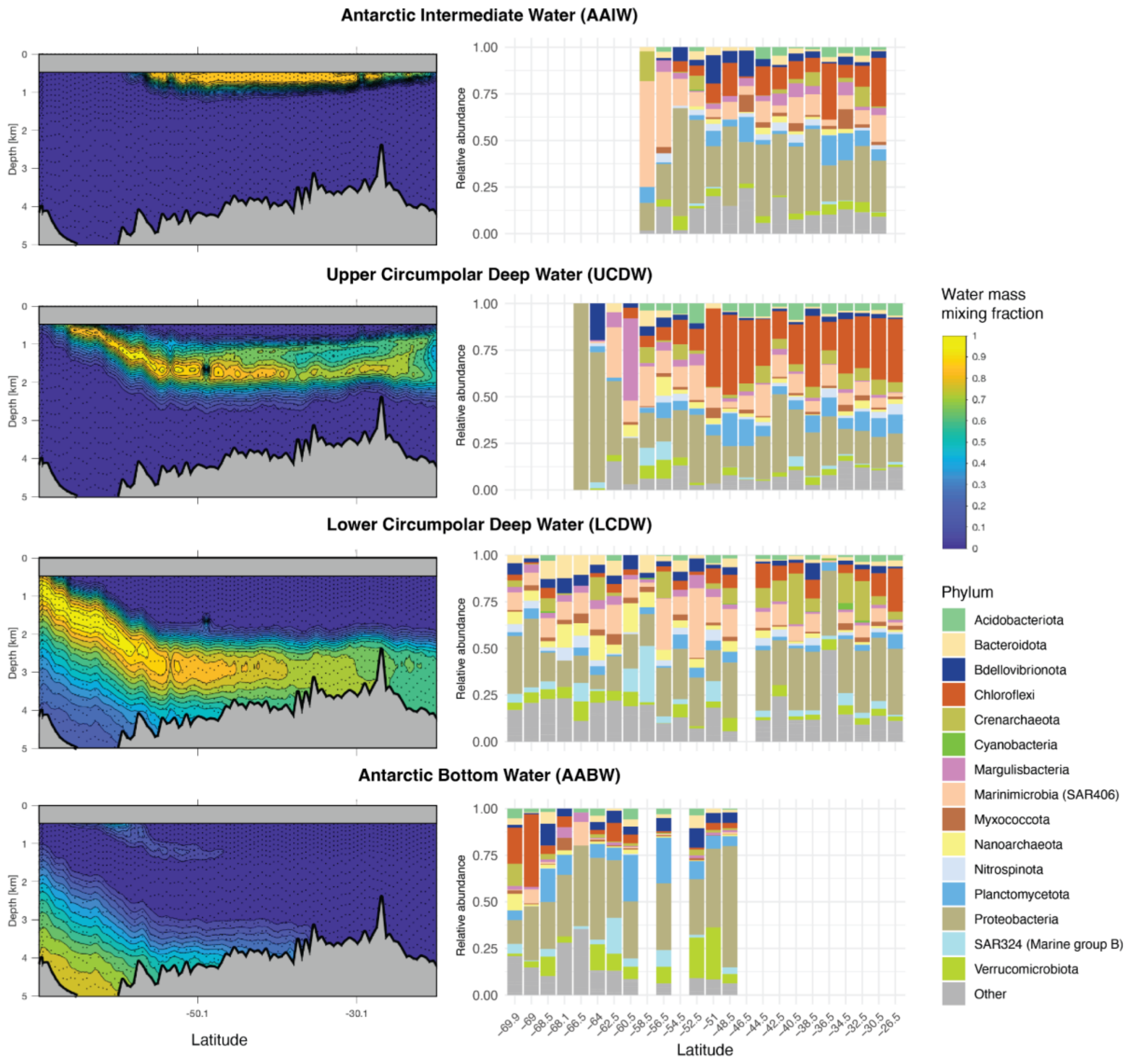


**Fig. S2. Taxonomic overview of ASVs endemic to each water mass.**

Left: section plots depicting the mixing fraction of water pertaining to dominant water masses (horizontal panels) as calculated via Optimum Multiparameter Analysis (OMP). The top 500 m of the water column (grey) was excluded due to vertical mixing and classified as “Upper water”. Right: taxonomic composition of 16S ASVs endemic to each water mass (detected only in samples with the highest mixing fraction pertaining to the water mass of interest).

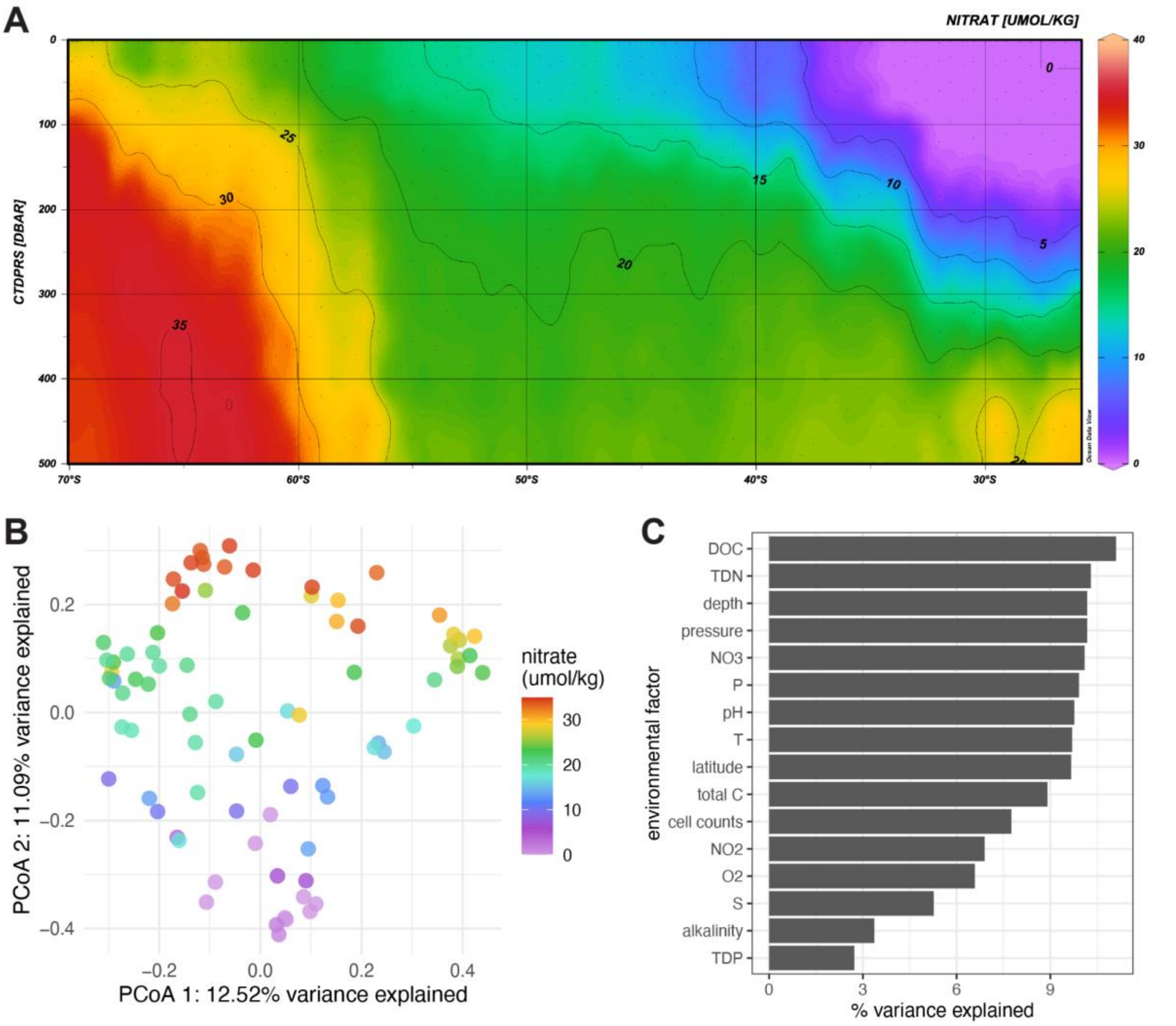


**Fig. S3. Nitrogen availability structures eukaryotic microbial communities.**

(A) Nitrate concentrations in the surface ocean. (B) Principal Coordinate Analysis (PCoA) plot of 18S rRNA amplicons in upper water (< 500m) colored by nitrate concentration. (C) Environmental factors structuring eukaryotic (18S rRNA) communities in upper water.

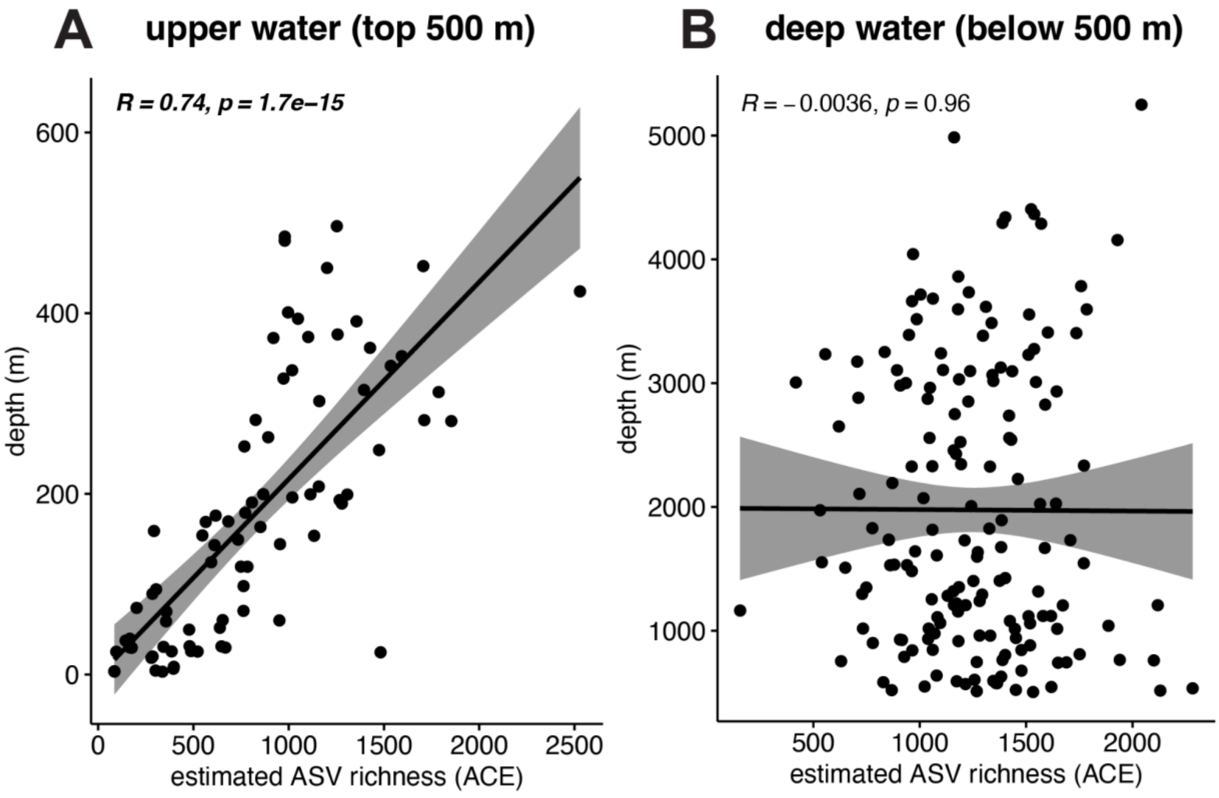


**Fig. S4. ASV richness is correlated with depth in upper water, but not deep water.**

Prokaryotic (16S rRNA amplicon) richness significantly increases with depth in upper water (A) but does not change significantly with depth below 500 m (B; Pearson correlation).

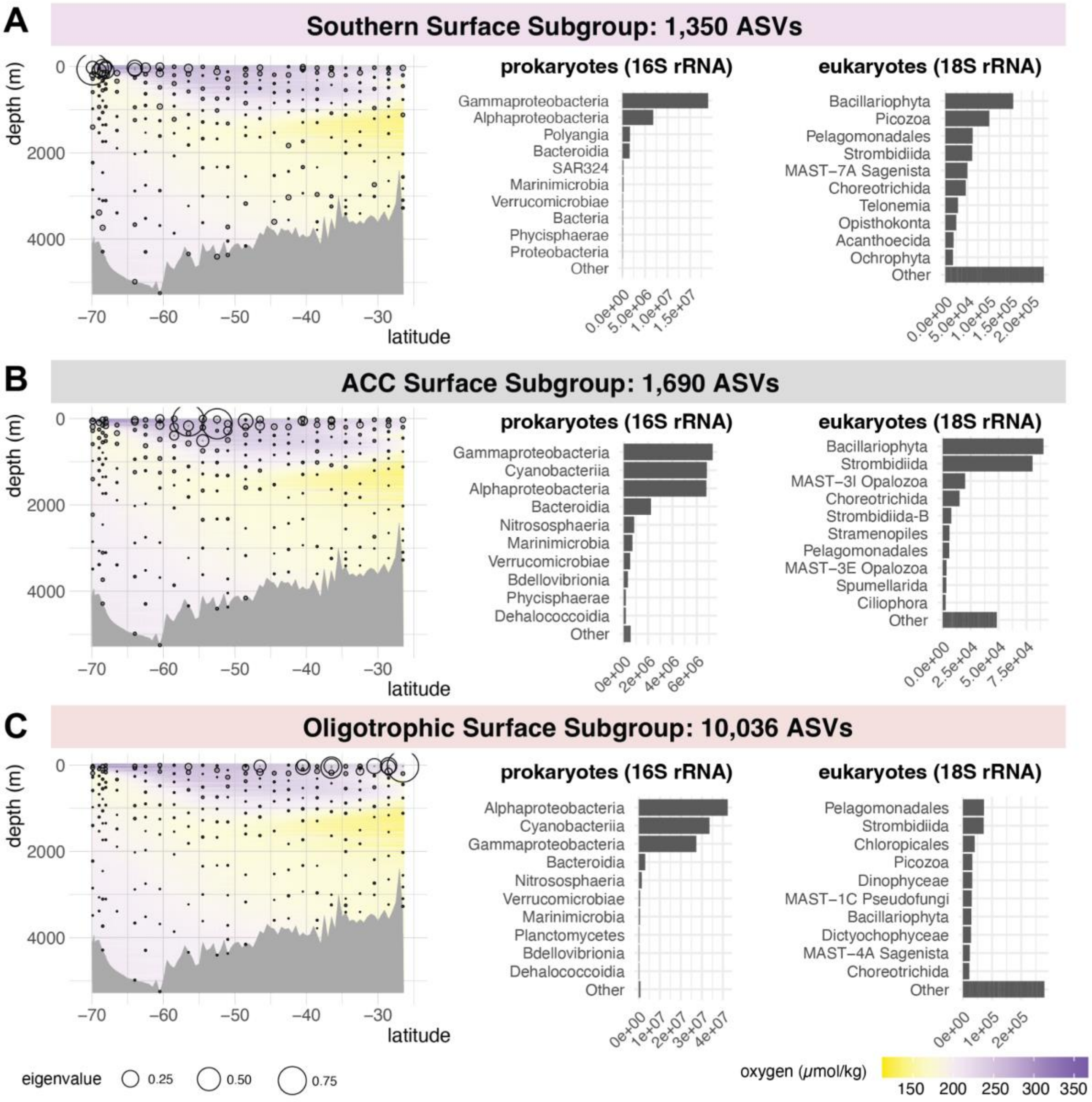


**Fig. S5. Surface ocean cohorts of co-occurring prokaryotic and eukaryotic ASV resolved through weighted group correlation network analysis (WGCNA).** Microbes self-assemble into a Southern Surface Subgroup (A), Antarctic Circumpolar Current (ACC) Surface Subgroup (B) and Oligotrophic Surface Subgroup (C). Left: section plots indicate the location of each microbial province across latitude and depth. Black circles indicate the average abundance profile for ASVs in the cohort. Right: bar graphs indicating the top 10 prokaryotic classes and eukaryotic orders present in the province, by estimated total amplicon copies/mL seawater filtered.

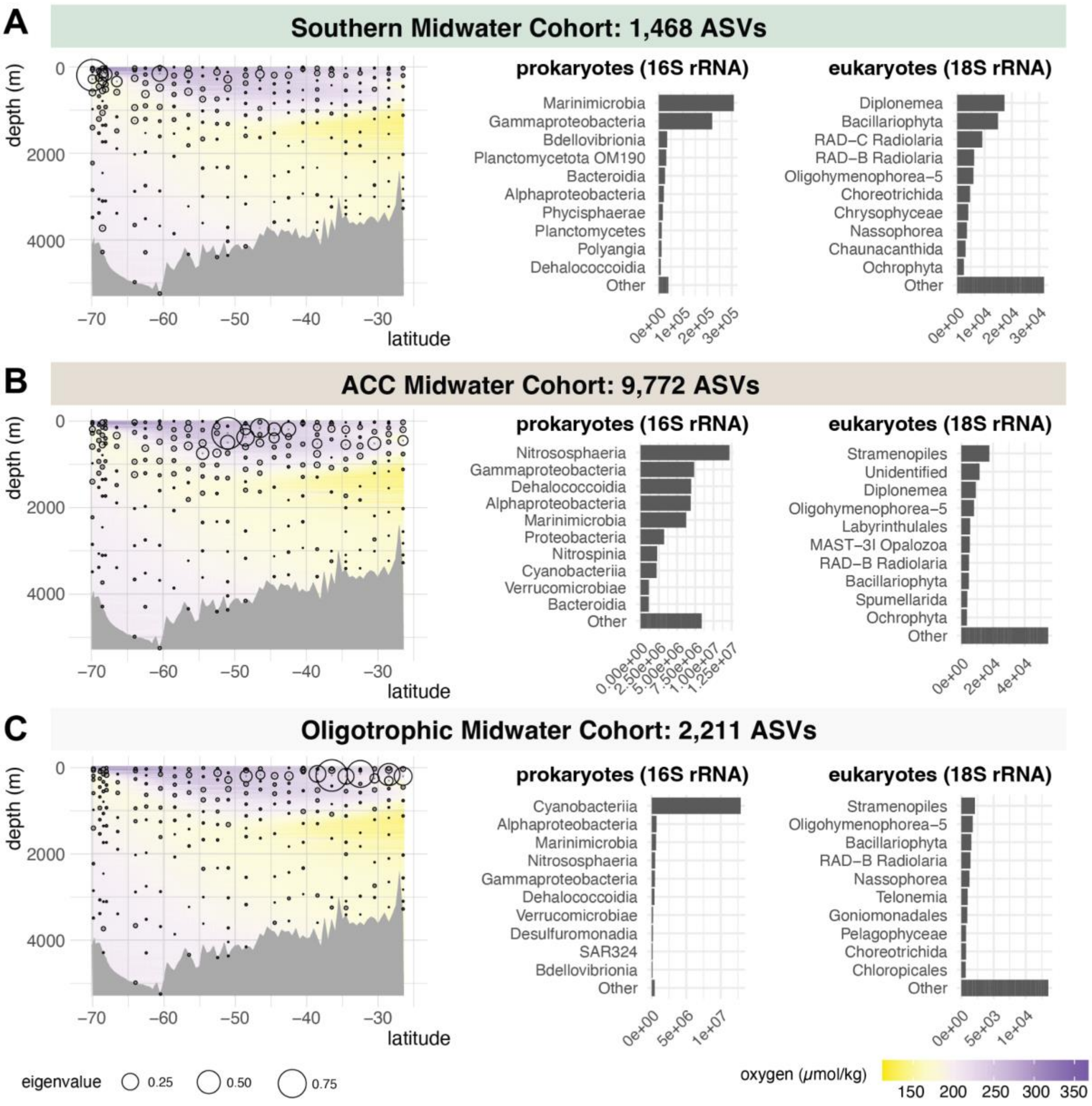


**Fig. S6. Midwater cohorts of co-occurring prokaryotic and eukaryotic ASV resolved through weighted group correlation network analysis (WGCNA).** Microbes self-assemble into a Southern Midwater Cohort (A), an Antarctic Circumpolar Current (ACC) Midwater Cohort (B) and an Oligotrophic Midwater Cohort (C). Left: section plots indicate the location of each microbial province across latitude and depth. Black circles indicate the average abundance profile for ASVs in the cohort. Right: bar graphs indicating the top 10 prokaryotic classes and eukaryotic orders present in the province, by estimated total amplicon copies/mL seawater filtered.

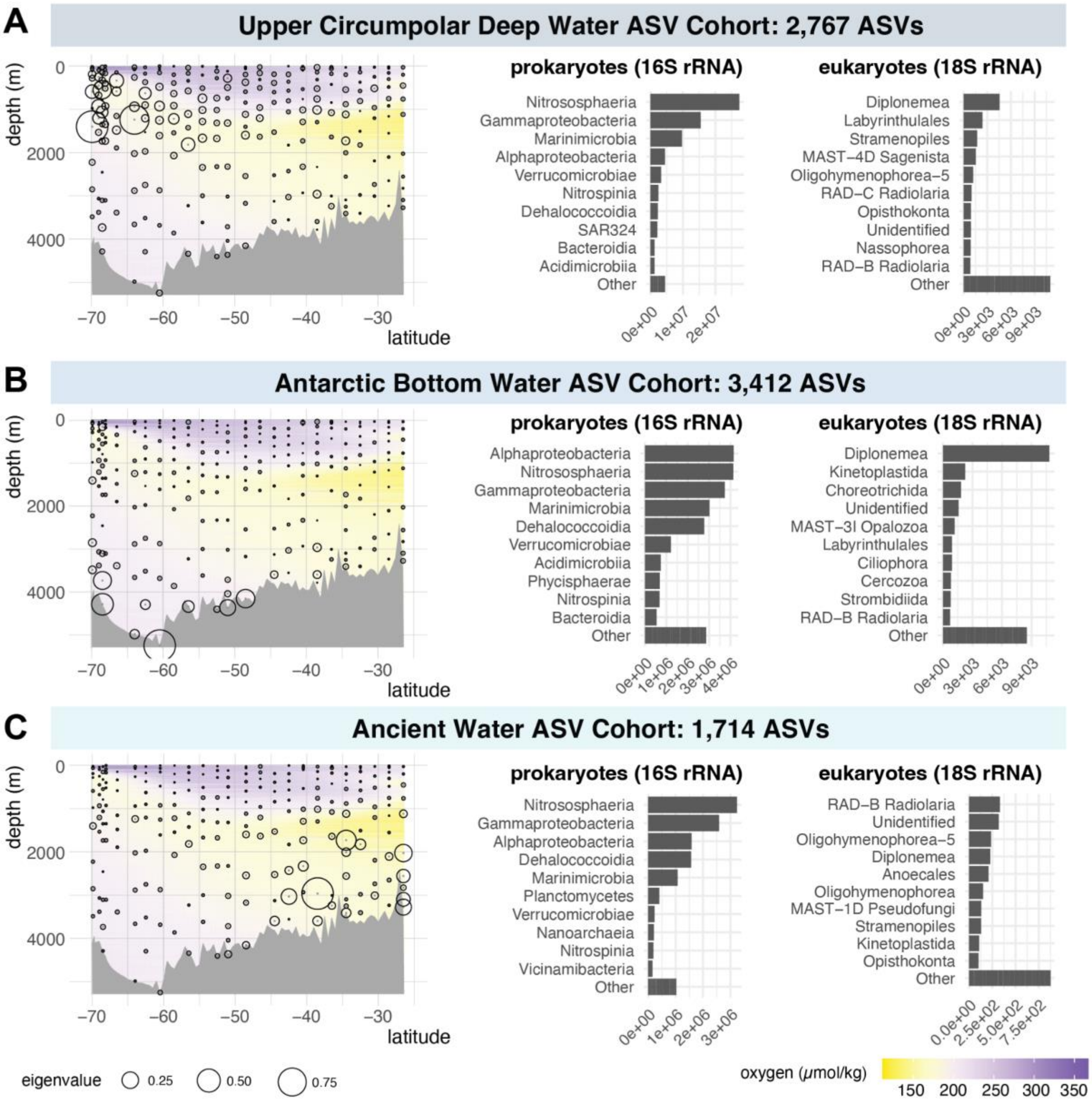


**Fig. S7. Deep ocean cohorts of co-occurring prokaryotic and eukaryotic ASV resolved through weighted group correlation network analysis (WGCNA).** ASVs recapitulate the Upper Circumpolar Deep Water (CDW) Cohort (A), Antarctic Bottom Water (AABW) Cohort (B) and Ancient Water Cohort (C) established with genomes. Left: section plots indicate the location of each microbial province across latitude and depth. Black circles indicate the average abundance profile for ASVs in the cohort. Right: bar graphs indicating the top 10 prokaryotic classes and eukaryotic orders present in the province, by estimated total amplicon copies/mL seawater filtered.

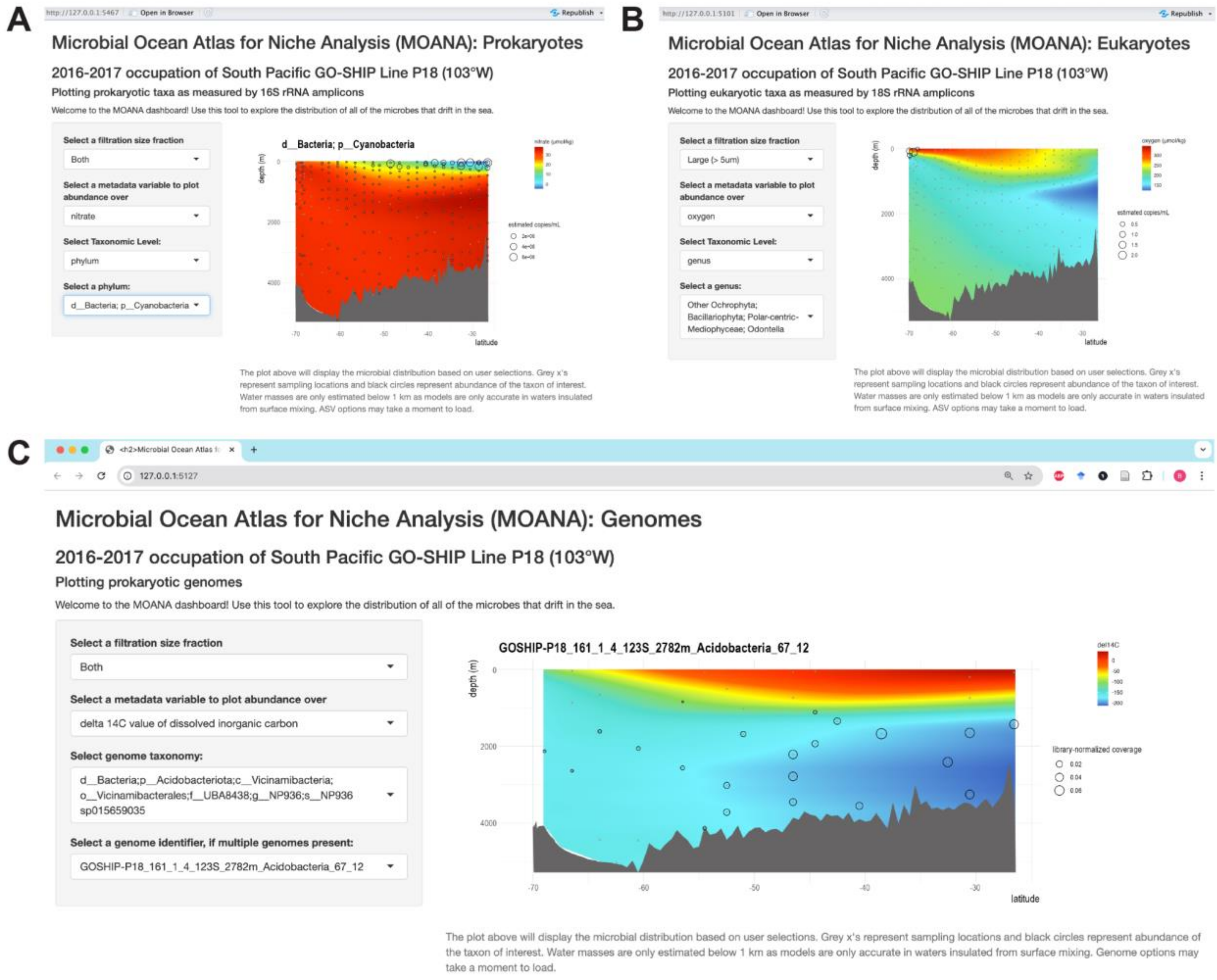


**Fig. S8. Example usage of the Microbial Ocean Atlas for Niche Analysis (MOANA) tool presented here.** The abundance of any microbe of interest, at taxonomic levels ranging from ASV to Phylum, can be plotted across latitude and depth resolved ocean sections of environmental variables that may explain their distribution. MOANA is available as a docker image at https://hub.docker.com/repositories/bkolody/ (see supplementary text for usage instructions). (A) Est. cyanobacterial 16S rRNA copies/mL plotted over a nitrate section using *MOANA: prokaryotes*, showing that they are most abundant in the nitrate-poor oligotrophic portion of the transect (as observed via cell counts, Fig. S16). (B) Est. 18S rRNA copies/mL of the diatom genus, *Odontella*, over a silicate section, plotted using *MOANA: Eukaryotes*, showing that it is most abundant in the Southern Zone where silicate is highest. (C) Library-normalized coverage of a *Vicinibacterales* genome over a delta 14C section, plotted using *MOANA: Genomes*, showing that it is most abundant in old water.

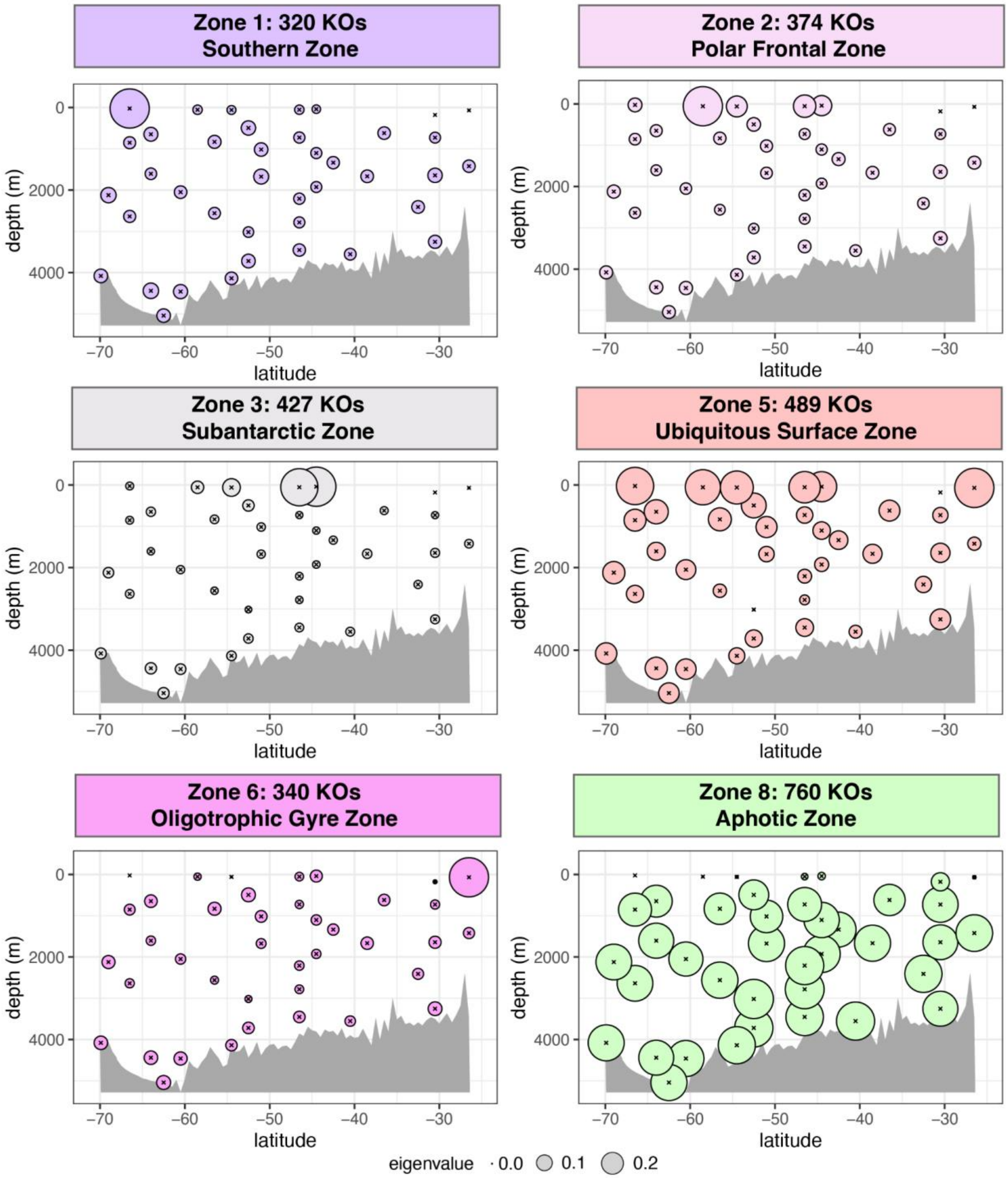


**Fig. S9. Microbial functional zones of the South Pacific (continued).** Section plots indicating average coverage pattern (eigensections) of the functional zones not pictured in the main text.

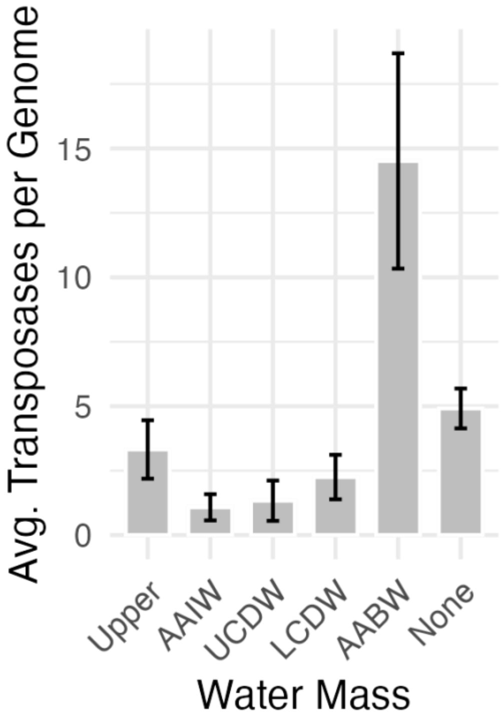


**Fig. S10. Genomes enriched in Antarctic Bottom Water (AABW) contain the most transposases.**

Average number of transposases present in each genome by water mass. “None” category represents genomes not significantly enriched in any given water mass (FDR> 0.05).

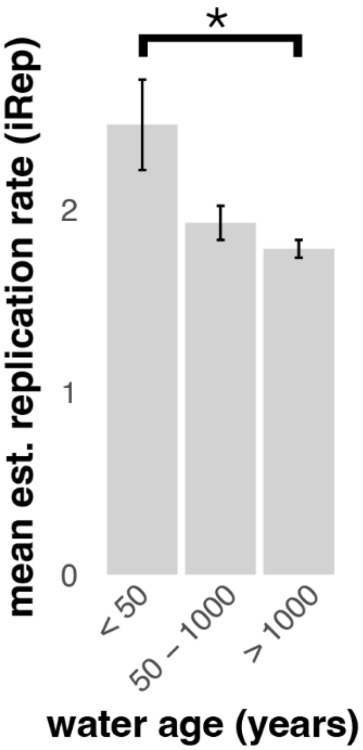


**Fig. S11. Genomes enriched in young water are predicted to replicate more quickly than those enriched in ancient water.**

Average index of replication (iRep) of genome bins significantly enriched (FDR < 0.05) in water < 50 years old, 50 -1000 years old, and > 1000 years old.

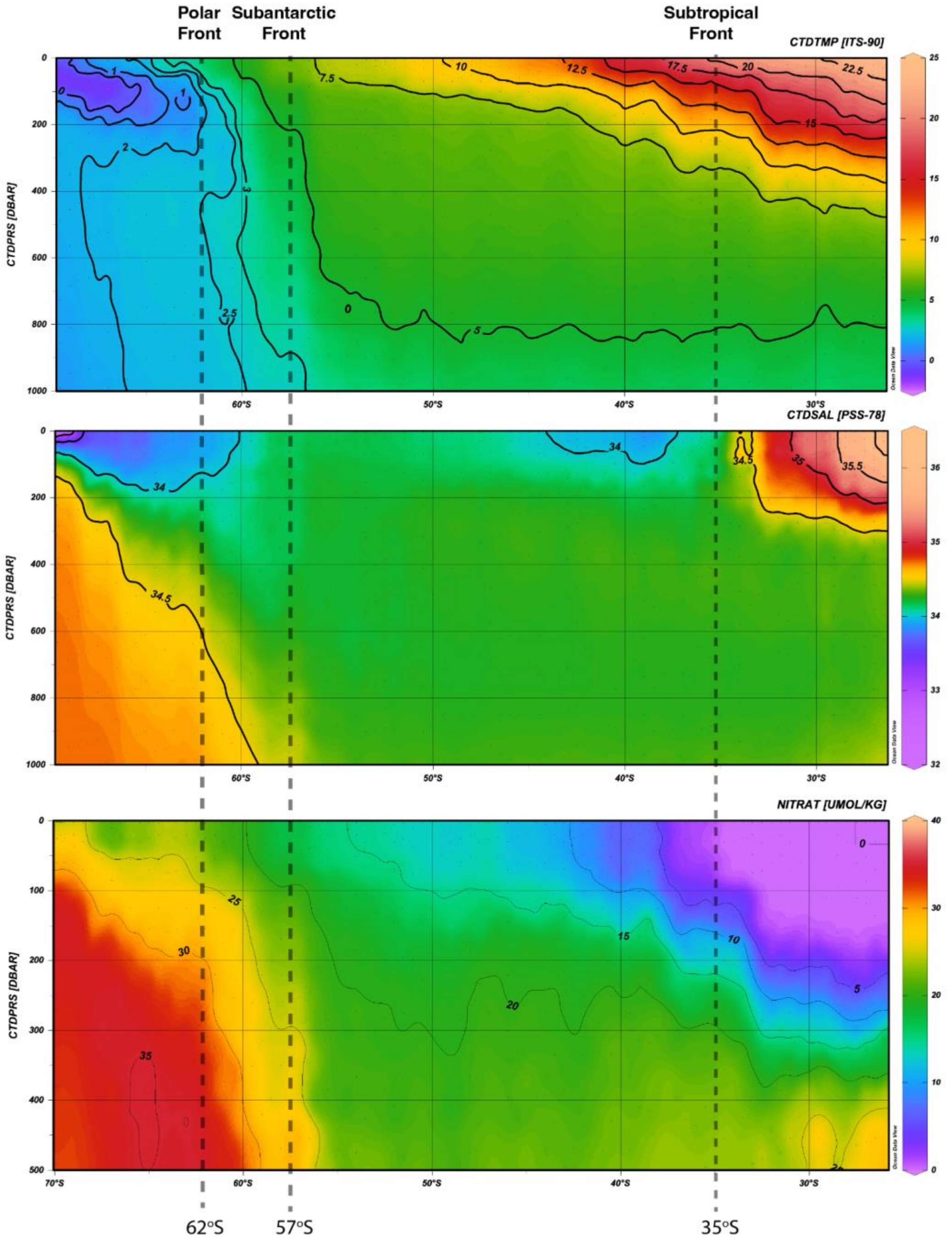


**Fig. S12. Visualization of the principal fronts defining major oceanographic zones of the P18 transect.** The Polar Front (PF) of the Antarctic Circumpolar Current (ACC) is defined as the northern limit of 2°C Antarctic Winter Water at 200 m deep. The Subantarctic Front (SAF) of the ACC is at 57°S at the northernmost limit of the low salinity intermediate layer. The Subtropical Front (STF), here at around 35°S, separates colder Subantarctic Water from the warmer, saltier, lower nutrient Subtropical Water of the oligotrophic gyre. Figure made in Ocean Data View.

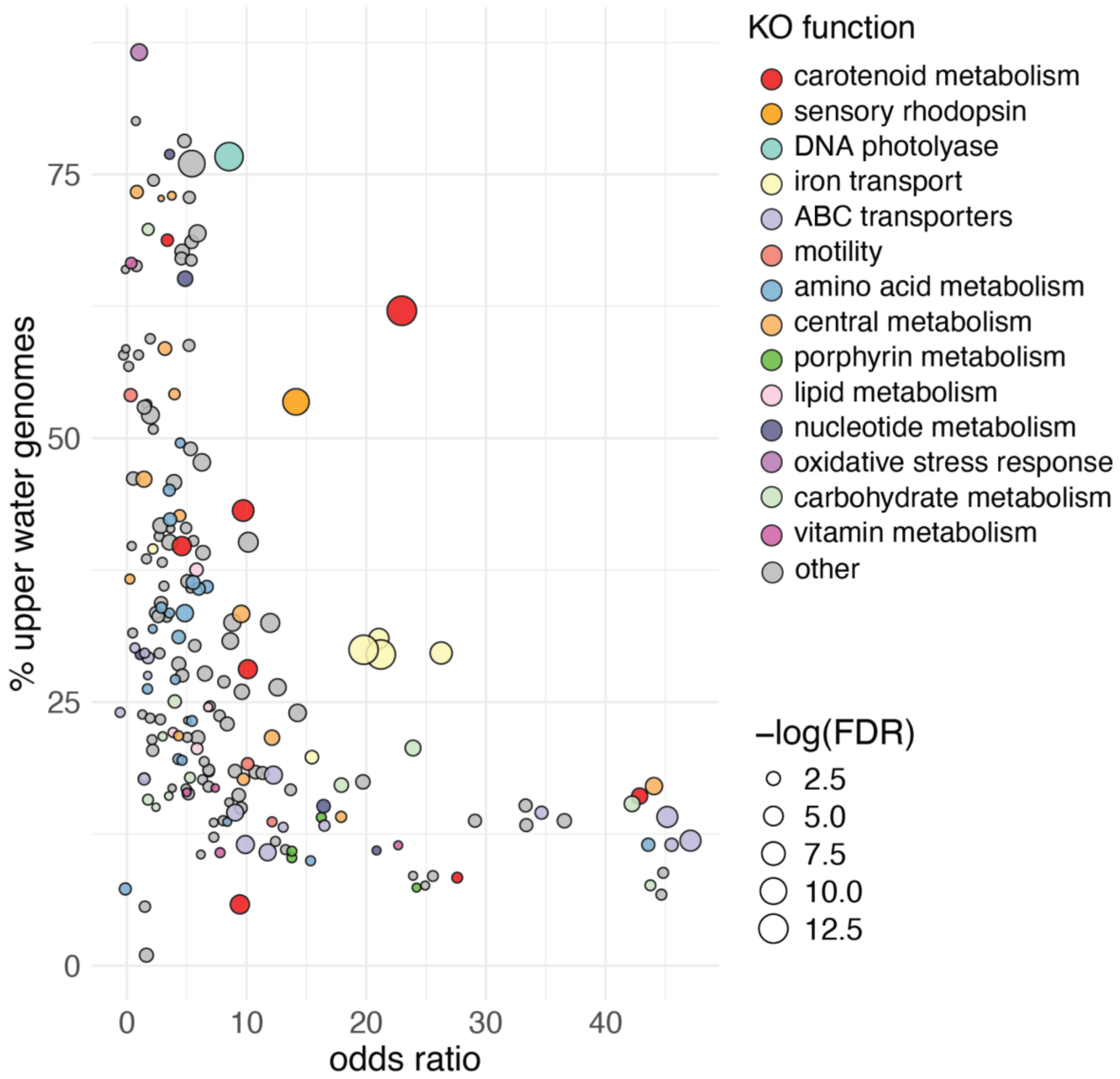


**Fig. S13. KEGG Orthology (KO) functions enriched Upper water genomes relative to all other genomes (Fisher FDR < 0.05).** Functions with infinite odds ratios were assigned the maximum non-infinite odds ratio for plotting. A jitter of 3 was added to both axes to disperse points for clearer viewing.

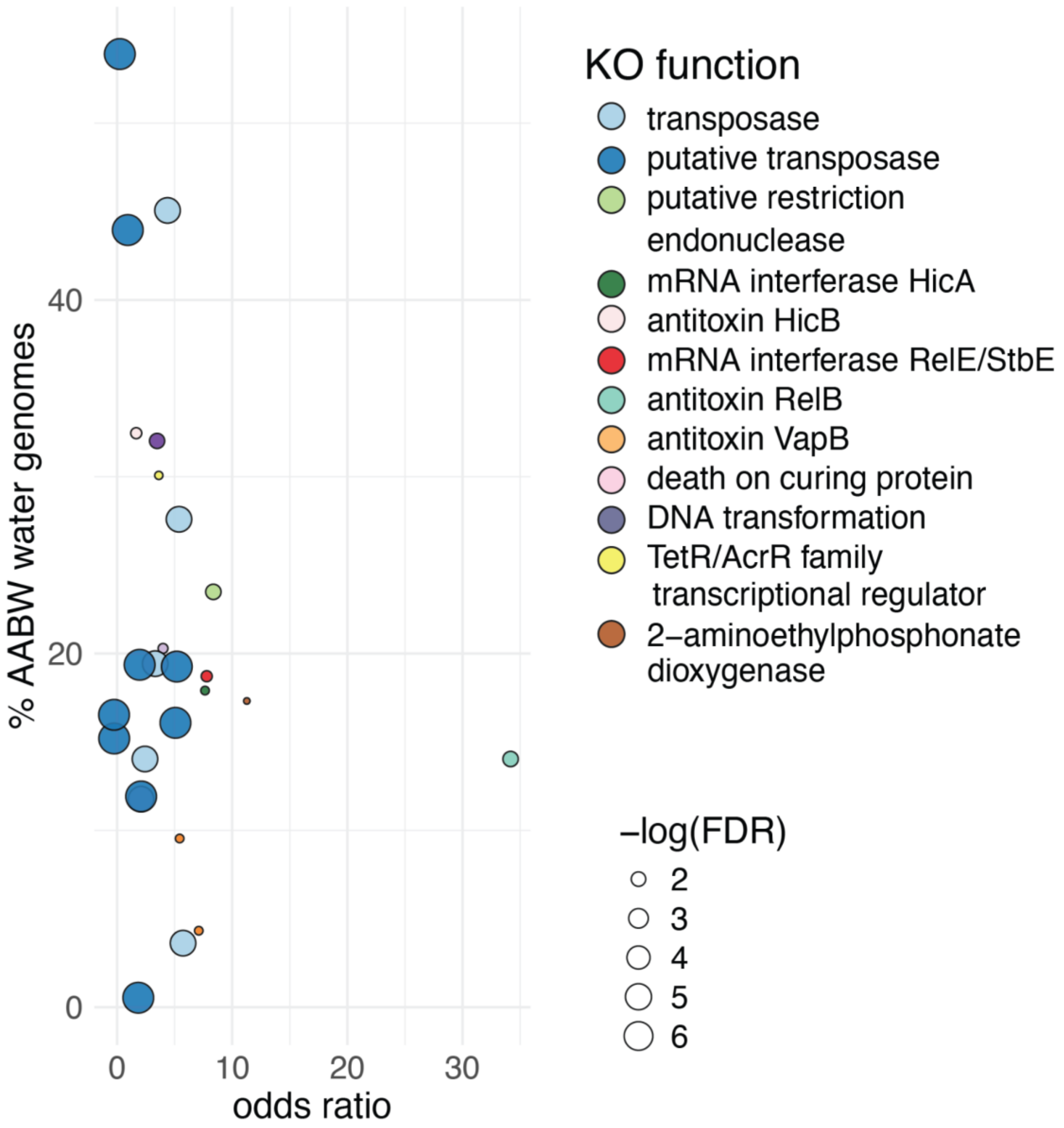


**Fig. S14. KEGG Orthology (KO) functions enriched Antarctic Bottom Water (AABW) genomes relative to all other genomes (Fisher FDR < 0.05).** A jitter of 3 was added to both axes to disperse points for clearer viewing.

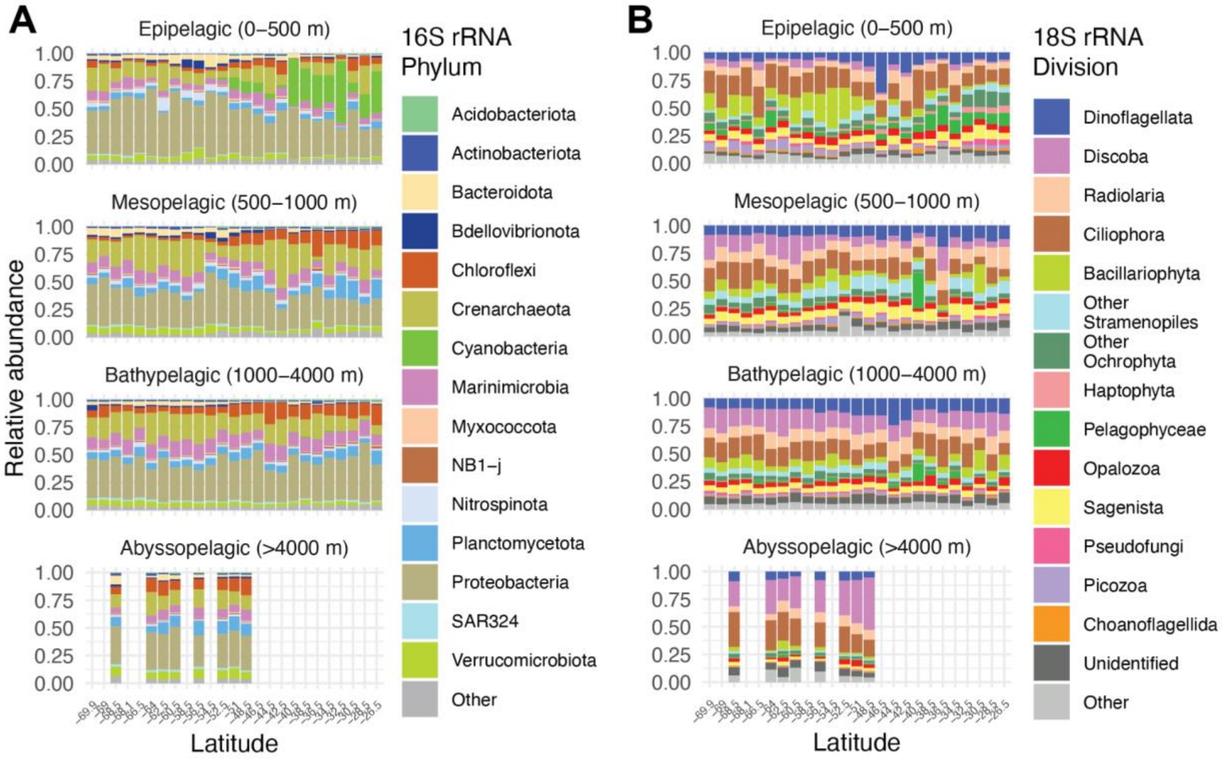


**Fig. S15. Overview of microbial community structure by latitude and depth.**
(A) Phylum-level community structure of 16S ASVs across latitude by ocean depth zone. (B) Division-level community structure of 18S ASVs across latitude by ocean depth zone.

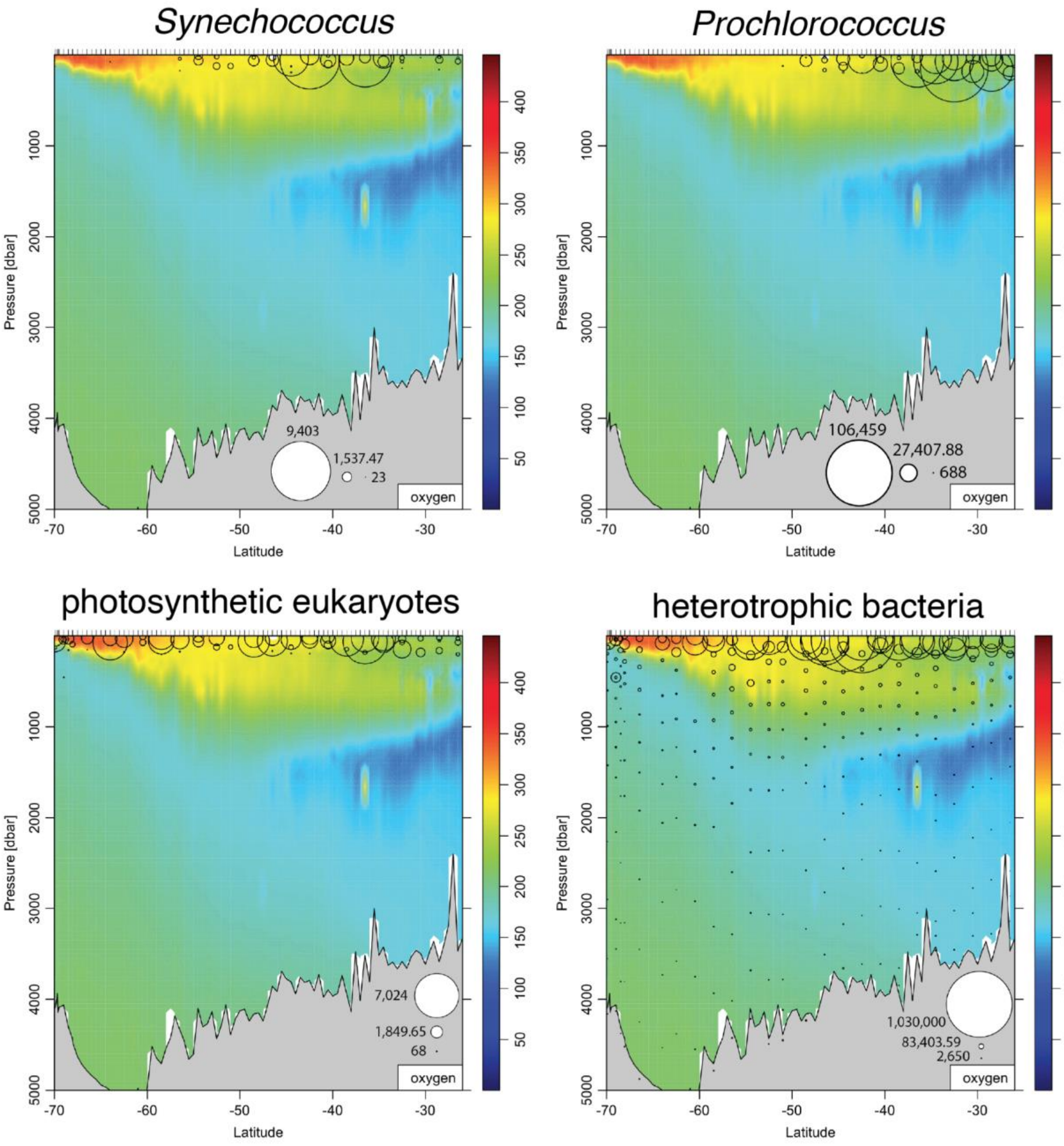


**Fig. S16. Cell counts.** Abundance of *Synechococcus* (top left), *Prochlorococcus* (top right), photosynthetic eukaryote (bottom left), and heterotrophic bacterial cells (bottom right) as measured by flow cytometry. Cell counts (open circles) are plotted on top of an oxygen section.

**Table S1. $R^2$ and FDR-adjusted p-values for PERMANOVAs testing the significance of environmental metadata in explaining variance of 16S rRNA and 18S rRNA amplicon community structure.**

| **Variable** | **16S $R^2$** | **16S FDR** | **18S $R^2$** | **18S FDR** |
|---|---|---|---|---|
| water mass | 0.22 | 0.001 | 0.124 | 0.001 |
| sample depth | 0.12 | 0.001 | 0.055 | 0.001 |
| CTD pressure | 0.12 | 0.001 | 0.054 | 0.001 |
| silicate | 0.117 | 0.001 | 0.06 | 0.001 |
| $SF_6$ | 0.114 | 0.001 | 0.072 | 0.001 |
| DELC14 | 0.11 | 0.001 | 0.063 | 0.001 |
| total dissolved nitrogen | 0.109 | 0.001 | 0.075 | 0.001 |
| alkalinity | 0.107 | 0.001 | 0.052 | 0.001 |
| CFC-12 | 0.106 | 0.001 | 0.062 | 0.001 |
| total carbon | 0.105 | 0.001 | 0.063 | 0.001 |
| CFC-11 | 0.102 | 0.001 | 0.06 | 0.001 |
| DELC13 | 0.097 | 0.001 | 0.057 | 0.001 |
| CTD oxygen | 0.088 | 0.001 | 0.055 | 0.001 |
| pH | 0.088 | 0.001 | 0.061 | 0.001 |
| dissolved organic carbon | 0.087 | 0.001 | 0.061 | 0.001 |
| nitrate | 0.087 | 0.001 | 0.06 | 0.001 |
| phosphate | 0.086 | 0.001 | 0.059 | 0.001 |
| bottle oxygen | 0.082 | 0.001 | 0.053 | 0.001 |
| heterotrophic bacteria cell counts | 0.082 | 0.001 | 0.054 | 0.001 |
| temperature | 0.072 | 0.001 | 0.051 | 0.001 |
| nitrite | 0.07 | 0.001 | 0.049 | 0.001 |
| photosynthetic eukaryotic cell counts | 0.066 | 0.001 | 0.048 | 0.001 |
| latitude | 0.066 | 0.001 | 0.047 | 0.001 |
| water age | 0.057 | 0.001 | 0.039 | 0.001 |
| salinity | 0.051 | 0.001 | 0.029 | 0.001 |
| total dissolved phosphorus | 0.044 | 0.001 | 0.028 | 0.001 |

| | | | | |
|---|---|---|---|---|
| Prochlorococcus cell counts | 0.038 | 0.001 | 0.031 | 0.001 |
| Synechococcus cell counts | 0.036 | 0.001 | 0.021 | 0.001 |
| total 18S rRNA copies/mL | 0.033 | 0.001 | 0.034 | 0.001 |
| total 16S rRNA copies/mL | 0.026 | 0.001 | 0.024 | 0.001 |
| $CCL_4$ | 0.011 | 0.006 | 0.007 | 0.011 |
| $N_2O$ | 0.011 | 0.004 | 0.007 | 0.008 |

**Table S2. Chao2 estimates of 16S rRNA diversity by water mass: Antarctic Bottom Water (AABW), Antarctic Intermediate Water (AAIW), Lower Circumpolar Deep Water (LCDW), Upper Circumpolar Deep Water (UCDW), and Upper Water (above 500 m). UCL (upper confidence limit) and LCL (lower confidence limit) represent the bounds of the confidence intervals for these diversity estimates.**

| Water Mass | Diversity | Observed | Estimator | s.e. | LCL | UCL |
|---|---|---|---|---|---|---|
| AABW | Species richness | 15664.334 | 314.642 | 15047.647 | 16281.021 | 15664.334 |
| AABW | Shannon diversity | 5310.396 | 47.641 | 5217.022 | 5403.771 | 5310.396 |
| AABW | Simpson diversity | 2801.072 | 17.112 | 2767.534 | 2834.611 | 2801.072 |
| AAIW | Species richness | 21078.356 | 271.89 | 20545.461 | 21611.251 | 21078.356 |
| AAIW | Shannon diversity | 7320.69 | 49.941 | 7222.807 | 7418.574 | 7320.69 |
| AAIW | Simpson diversity | 3866.729 | 21.682 | 3824.234 | 3909.225 | 3866.729 |
| LCDW | Species richness | 34452.673 | 408.759 | 33651.52 | 35253.825 | 34452.673 |
| LCDW | Shannon diversity | 7480.429 | 37.309 | 7407.304 | 7553.553 | 7480.429 |
| LCDW | Simpson diversity | 3540.793 | 13.312 | 3514.702 | 3566.884 | 3540.793 |
| UCDW | Species richness | 24043.511 | 311.935 | 23432.13 | 24654.893 | 24043.511 |
| UCDW | Shannon diversity | 6878.991 | 38.462 | 6803.607 | 6954.374 | 6878.991 |
| UCDW | Simpson diversity | 3579.979 | 16.885 | 3546.886 | 3613.073 | 3579.979 |
| Upper | Species richness | 35009.985 | 360.52 | 34303.379 | 35716.592 | 35009.985 |
| Upper | Shannon diversity | 11997.151 | 68.621 | 11862.656 | 12131.646 | 11997.151 |
| Upper | Simpson diversity | 5360.277 | 30.38 | 5300.734 | 5419.821 | 5360.277 |

**Table S3. Chao2 estimates of 18S rRNA diversity by water mass: Antarctic Bottom Water (AABW), Antarctic Intermediate Water (AAIW), Lower Circumpolar Deep Water (LCDW), Upper Circumpolar Deep Water (UCDW), and Upper Water (above 500 m). UCL (upper confidence limit) and LCL (lower confidence limit) represent the bounds of the confidence intervals for these diversity estimates.**

| Assemblage | Diversity | Observed | Estimator | s.e. | LCL | UCL |
|---|---|---|---|---|---|---|
| AABW | Species richness | 9948 | 17310.798 | 205.949 | 16907.146 | 17714.449 |
| AABW | Shannon diversity | 5947.396 | 7482.93 | 44.511 | 7395.69 | 7570.171 |
| AABW | Simpson diversity | 4106.023 | 4388.608 | 23.149 | 4343.237 | 4433.98 |
| AAIW | Species richness | 12398 | 21147.073 | 217.131 | 20721.504 | 21572.642 |
| AAIW | Shannon diversity | 6920.727 | 8422.454 | 46.986 | 8330.363 | 8514.544 |
| AAIW | Simpson diversity | 4653.314 | 4904.478 | 23.833 | 4857.766 | 4951.19 |
| LCDW | Species richness | 15528 | 27994.732 | 324.114 | 27359.48 | 28629.984 |
| LCDW | Shannon diversity | 6456.475 | 7309.877 | 27.339 | 7256.294 | 7363.46 |
| LCDW | Simpson diversity | 3965.321 | 4049.979 | 12.993 | 4024.514 | 4075.444 |
| UCDW | Species richness | 14418 | 24320.367 | 238.709 | 23852.505 | 24788.229 |
| UCDW | Shannon diversity | 6661.793 | 7632.033 | 27.649 | 7577.842 | 7686.224 |
| UCDW | Simpson diversity | 4216.21 | 4336.453 | 14.504 | 4308.025 | 4364.881 |
| Upper | Species richness | 30918 | 60274.906 | 527.439 | 59241.146 | 61308.667 |
| Upper | Shannon diversity | 11837.632 | 13902.073 | 43.501 | 13816.813 | 13987.333 |
| Upper | Simpson diversity | 6459.106 | 6609.418 | 15.604 | 6578.835 | 6640.001 |

**Table S4. 16S rRNA ASVs closely related to (> 99% ID over > 200 bp) experimentally validated psychrophiles (P) and psychropiezophiles (PP) described in Lauro et. al 2006.**

| ASV | cohort | Genbank accession | % ID | alignment length | extremophile hit | type |
|---|---|---|---|---|---|---|
| 844e54326efc6ea7e5c7d1ae9b30f49a | ACC Midwater Cohort | AB038032 | 99.73 | 373 | Photobacterium sp. HAR72 | P |
| 8c21844441c36ccd50e40d83874ef72e | ACC Midwater Cohort | AB038033 | 100 | 373 | Moritella marina ATCC15381 | P |
| b187b7d474cc712409bffc2fcb24d080 | Ancient Water Cohort | D21225 | 99.46 | 373 | Shewanella violacea DSS12 | PP |
| 314b1f3a4d181cad945ab13f0f1d80ef | AABW Cohort | AJ387905 | 99.73 | 374 | Carnobacterium gallinarum DSM4847 | P |
| 347b9a4ac1159fe68974c214a3be663e | AABW Cohort | AB023378 | 99.73 | 373 | Psychromonas marina 4-22 | P |
| 3ae15dea4f26e52fcb2f8544748fc11e | AABW Cohort | AB023378 | 100 | 373 | Psychromonas marina 4-22 | P |
| 97034b5f8ca9938ced6fa850ac7e6b05 | AABW Cohort | AB002630 | 100 | 373 | Colwellia maris ABE-1 | P |
| cb10fe8f1f6c1004afc1ab1da25cd00b | AABW Cohort | AB002630 | 99.73 | 373 | Colwellia maris ABE-1 | P |
| d59d0e589695a463c5ea5cbd5bc1fb72 | AABW Cohort | AF396670 | 100 | 373 | Colwellia psychrerythraea 34H | P |
| 194880ce683f9cee38d1aa68ac9f6d77 | AABW Cohort | AB094413 | 100 | 373 | Psychromonas hadaliensis K41G | PP |
| 52a4e081a2aa43f6250875a730f703ed | AABW Cohort | AB094413 | 99.73 | 373 | Psychromonas hadaliensis K41G | PP |
| f25d31bc417e8054ee4036ffd7066e41 | AABW Cohort | D21225 | 99.73 | 373 | Shewanella violacea DSS12 | PP |
| 050ccc9a297a459818a1224bdeed4d6a | Oligotrophic Surface Subgroup | AB038033 | 99.46 | 373 | Moritella marina ATCC15381 | P |
| a650cc7e570c35c463b2a5af94c9e0ef | Oligotrophic Surface Subgroup | AB038033 | 99.73 | 373 | Moritella marina ATCC15381 | P |

| | | | | | | |
|---|---|---|---|---|---|---|
| c22de5a7b214ad0ccb102c8d5b723e15 | Oligotrophic Surface Subgroup | AB038031 | 100 | 373 | Photobacterium sp. HAR23 | P |
| b9857731cc0762731ea5e6feb996cfec | UCDW Cohort | AB038033 | 99.73 | 373 | Moritella marina ATCC15381 | P |
| a5a8bd52a9d6ba2db04c65c320a11a7c | UCDW Cohort | D21225 | 99.2 | 373 | Shewanella violacea DSS12 | PP |
| 03fdefd49147b542fcc189d68d06f220 | NA | AF530149 | 99.2 | 373 | Shewanella gelidimarina MGP-71 | P |
| 115a27fb6f82f52571e21b84083f4925 | NA | AF396670 | 100 | 218 | Colwellia psychrerythraea 34H | P |
| 1bce49d4982bd52d447b644db8d756c9 | NA | AB083406 | 99.73 | 374 | Marinilactibacillus psychrotolerans M13-2 | P |
| 417be7f9150810a26da7ce11bbad000e | NA | AB083406 | 100 | 374 | Marinilactibacillus psychrotolerans M13-2 | P |
| 6fffed5e90af3e9dbb91843491a5cf61 | NA | AB038031 | 99.46 | 373 | Photobacterium sp. HAR23 | P |
| 73218d218e1d383df00e0f9afe3754aa | NA | AB023378 | 99.73 | 373 | Psychromonas marina 4-22 | P |
| 7f34b641ec728251f961338a0485a96a | NA | AB038032 | 99.46 | 373 | Photobacterium sp. HAR72 | P |
| 8041a5cfb8169566fa3ba16274daeb3f | NA | AB002630 | 99.46 | 373 | Colwellia maris ABE-1 | P |
| 875e7567d2684880101cb40b3552cbe9 | NA | AB023378 | 99.73 | 373 | Psychromonas marina 4-22 | P |
| a550a3c0d3b6fc3b8d6424835e16a60a | NA | AB038033 | 99.2 | 373 | Moritella marina ATCC15381 | P |
| a614edfc838f8d7abdb52c82573680b7 | NA | AB023378 | 99.73 | 373 | Psychromonas marina 4-22 | P |
| c37e8fab12251eae82353be6d9556661 | NA | AB023378 | 99.73 | 373 | Psychromonas marina 4-22 | P |

| | | | | | | |
|---|---|---|---|---|---|---|
| d531d6c5112b6c6709fa789129713248 | NA | AJ387905 | 99.19 | 371 | Carnobacterium gallinarum DSM4847 | P |
| e3ed124c6cf9fdcf5435cf53e8c3db9b | NA | AB038033 | 99.73 | 373 | Moritella marina ATCC15381 | P |
| fd04a0453e5285ccaa021179e9391c8b | NA | AB002630 | 99.46 | 373 | Colwellia maris ABE-1 | P |
| 19222ed044e50b5220153a4f39c68175 | NA | AB094413 | 99.46 | 373 | Psychromonas hadaliensis K41G | PP |
| 19b5a7cd571674a307715773f15ece1a | NA | D21225 | 100 | 373 | Shewanella violacea DSS12 | PP |
| 6db8635003df6c1d27308a0c75e92224 | NA | D21225 | 99.73 | 373 | Shewanella violacea DSS12 | PP |
| 6f503bd791abedea346294fab609f2dd | NA | AB003191 | 99.46 | 373 | Photobacterium profundum SS9 | PP |
| 8725dd9a4be31e0803e779e6b871f62f | NA | AB008797 | 100 | 217 | Moritella yayanosii DB21MT-5 | PP |
| 9a2aa82de9a178a6ba67f9f70e0dfb2b | NA | D21225 | 99.2 | 373 | Shewanella violacea DSS12 | PP |

**Table S5. Source water definitions for determining the mixing fraction of water masses of the South Pacific.**

| Water mass | Full name | Lat. | Lon. | depth (m) | $\sigma_\theta$ (kg/m$^3$) | potential T (°C) | salinity (PSS-78) | oxygen (µmol/kg) | PO4 (µmol/kg) | NO3 (µmol/kg) | Si (µmol / kg) |
|---|---|---|---|---|---|---|---|---|---|---|---|
| AABW | Antarctic bottom water | 67°S | 30° W | 4750 | 27.8584 | -0.8793 | 34.64 | 6.041 | 2.21 | 31.3 | 117 |
| NPIW | North Pacific Intermediate Water | 37°N | 160° E | 471.6545 | 26.8 | 5.498067 | 33.972 | 3.45309414 | 2.038 | 29.23 | 62.23 |
| LCDW | Lower Circumpolar Deep Water | 55°S | 150° W | 2250 | 27.80005 | 1.46716 | 34.74 | 4.473 | 2.24 | 32.4 | 103 |
| UCDW | Upper Circumpolar Deep Water | 53°S | 150° W | 1500 | 27.6134 | 2.35106 | 34.59 | 4.016 | 2.39 | 34.5 | 79.6 |
| AAIW | Antarctic Intermediate Water | 55°S | 85° W | 635.1783 | 27.1 | 4.626677 | 34.221 | 5.88961522 | 1.854 | 27.14 | 17.62 |
| PDW | Pacific Deep Water | 35°N | 150° W | 2250 | 27.69751 | 1.65573 | 34.62 | 1.864 | 2.91 | 42.3 | 176 |

**Table S6. Genomic context of scaffolds containing death-on-curing (DOC) proteins including toxin-antitoxin systems (TASs).**

| **Scaffold (genome taxonomy)** | **DOC gene position** | **Best TAS annotation** | **TAS gene position** | **Genomic context (gene position, description)** |
|---|---|---|---|---|
| GOSHIP-P18_121_1_9_6S_1421m_2_2509 (*Sulfitobacter* sp001634775) | 7 | type II TAS: Phd/YefM family | 6 | DNA-binding protein (1), MATE family efflux transporter (4, oxidative stress and salt stress response) |
| GOSHIP-P18_129_2_8_29S_1643m_2_4109 (Dehalococcoidia UBA11650 sp002710215) | 3 | Antitoxin component of the MazEF TAS, type II Phd/YefM family TAS, type II TAS VapC family toxin | 4,6,7 | Fur family ferric uptake regulator (1) |
| GOSHIP-P18_149_1_2_86S_3553m_2_2257 (Dehalococcoidia Io17-Chloro-G2) | 16 | none identified | NA | class III cytochrome c family protein (1, possibly involved in anaerobic iron respiration) |
| GOSHIP-P18_149_1_2_86S_3553m_1589 (Hydrogenedentiales DTSJ01) | 8 | Antitoxin component of MazEF TAS | 9 | Clp protease (7, involved in fatty-acid metabolism) |
| GOSHIP-P18_170_1_9_151S_1674m_66 (*Alteromonas* sp016405965) | 8 | type II TAS HipA family toxin | 29 | phage integrases (6, 7), transposases (12, 13, 18) |
| GOSHIP-P18_193_1_1_218L_5041m_15844 (Moraxellaceae CAJXOR01) | 1 | none identified | NA | formite/nitrite transporter (3), Cobalamin-independent methionine synthase (4) |

| | | | | |
|---|---|---|---|---|
| GOSHIP-P18_193_1_1_218L_5041m_49000 (Rhodospirillales Casp-alpha2) | 2 | Antitoxin component of MazEF TAS | 1 | RhaT transporter (3), C4 dicarboxylatetransporter permease (4) |
| GOSHIP-P18_193_1_1_218S_5041m_2_10065 (Alphaproteobacteria JAAXGF01) | 6 | Antitoxin component of MazEF TAS | 5 | serine/threonine dehydratase (4, amino acid catabolism), enoyl-CoA hydratase (1, fatty acid metabolism) |
| GOSHIP-P18_193_1_1_218S_5041m_19 (JAAXHH01 phylum, CAJXCW01 sp913051705) | 180 | type II TAS PemK/MazEF family toxin (175), Antitoxin component of MazEF TAS | 175, 179 | Plasmid maintenance system antidote protein VapI (169) |
| GOSHIP-P18_193_1_1_218S_5041m_5813 (JAAXHH01 phylum, CAJXCW01 sp913051705) | 1 | none identified | NA | xylose isomerase (6, hemicellulose degradation), Amidohydrolase 2 (3, break down of peptides, nucleotides, and other organic compounds containing amide bonds) |
| GOSHIP-P18_193_1_1_218S_5041m_1845 (Pedosphaerales GCA-2715965) | 6 | Antitoxin component of MazEF TAS | 7 | Glucose/arabinose dehydrogenase (3), Dabb family protein (11, salt stress response) |
| GOSHIP-P18_193_1_1_218S_5041m_97 (Longimicrobiales UBA1138) | 130 | MazEF family AbrB-like protein | 131 | M15 family metallopeptidase (132) |

| | | | | |
|---|---|---|---|---|
| GOSHIP-P18_193_1_1_218S_5041m_126 (Pseudomonadales UBA9659) | 11 | addiction module antidote | 12 | Fic (Filamentation induced by cAMP) family protein (10; morphological stress response to DNA damage or nutrient limitation), integrase (15), DNA uptake protein comEA (16), LapA family protein (18, biofilm formation), integration host factor subunit beta (19, TF involved in stress response) |
| GOSHIP-P18_196_1_3_231S_4440m_2_3249 (*Psychrobacter* sp000511655) | 9 | none identified | NA | LysR (7, environmental stress response), transposase (10) |
| GOSHIP-P18_196_1_3_231S_4440m_2_780 (Pseudohongiellaceae UBA9145 sp913054495) | 2 | none identified | NA | nitrogen regulatory protein PII (12), ammonium transporter (13), cold-shock protein (25) |
| GOSHIP-P18_196_1_3_231S_4440m_932 (Pedosphaerales UBA1096) | 32 | none identified | NA | tonB iron transporter (1), sulfatase (13, 14), drug/metabolite transporter (29) |
| GOSHIP-P18_210_1_1_290L_4079m_2_1576 (SAR324 Arctic96AD-7 sp016764995) | 11 | Antitoxin component of MazEF TAS | 10 | integrases (3,4), phage integrase (5) |

**Data S1. (separate file)**

Abundance and metadata associated with curated marine genomes (n = 307 prokaryotic MAGs). This spreadsheet contains genome names, quality and completeness metrics from checkM, taxonomic annotations from GTDB-Tk, WGCNA cohort pertinence, statistical enrichment in across water masses and water age categories, iRep replication rate estimates, and mean coverage values from mapping across all samples. iRep "NA" values indicate that the iRep value could not be calculated with confidence for the given genome. Genome size is given in bp. Genome names are composed of the following fields, separated by underscores: library name, ggkbase taxonomic annotation, approximate % GC, and approximate genome coverage in the library the genome was isolated from. Library names have the following fields, separated by underscores: cruise line, cruise station, CTD cast number, niskin number, molecular sample identifier, size class from serial filtration (L= large, 0.5 um filter; S = small, 0.22 um filter), and collection depth in meters. Libraries from the second sequencing effort are appended with "_2".

**Data S2. (separate file)**

16S rRNA amplicon sequence variant (ASV) counts with SILVA 138 taxonomic annotations and cohort affiliations. This is a large file best viewed in R or similar rather than excel.

**Data S3. (separate file)**

18S rRNA amplicon sequence variant (ASV) counts with PR2 taxonomic annotations and cohort affiliations. This is a large file best viewed in R or similar rather than excel.

**Data S4 (separate file)**

Environmental metadata associated with molecular samples. Units are as follows: depth (meters); pressure (dbar); temperature (ITS-90); salinity (PSS-78); oxygen, silicate, nitrate, nitrite, phosphate, total carbon, alkalinity, dissolved organic carbon, total dissolved nitrogen, total dissolved phosphorus (μmol/kg); CFCs, $CCl_4$ (pmol/kg); $N_2O$ (nmol/kg); $SF_6$ (fmol/kg); $\delta^{13}C$, $\delta^{14}C$ (per mille).

**Data S5. (separate file)**

Number and percentage of 16S rRNA and 18S rRNA ASVs unique to each sample. Missing 18S rRNA values correspond to samples for which 16S samples passed quality control thresholds but 18S rRNA samples did not (see Methods).

**Data S6. (separate file)**

KEGG Orthologies (KOs) pertaining to each functional zone, along with the number of genomes they are found in and total library normalized coverage they represent. This is a large file best viewed in R or similar rather than excel.

**Data S7. (separate file)**

Enrichment and depletion of KEGG Orthologies (KOs) from eukaryotic scaffolds in each water mass: Antarctic Bottom Water (AABW), Antarctic Intermediate Water (AAIW), Lower Circumpolar Deep Water (LCDW), Upper Circumpolar Deep Water (UCDW), and Upper Water (above 500 m).

**Data S8. (separate file)**

Statistical enrichment and depletion of KEGG Orthology (KO) annotation of genes across water masses based on relative genome coverage. Water mass acronyms are as follows: Antarctic Bottom Water (AABW), Antarctic Intermediate Water (AAIW), Lower Circumpolar Deep Water (LCDW), Upper Circumpolar Deep Water (UCDW), and Upper Water (above 500 m).

**Data S9. (separate file)**

Enrichment of Carbohydrate-Active enZYmes (CAZys) across water masses: Antarctic Bottom Water (AABW), Antarctic Intermediate Water (AAIW), Lower Circumpolar Deep Water (LCDW), Upper Circumpolar Deep Water (UCDW), and Upper Water (above 500 m). This is a large file best viewed in R or similar rather than excel.
